# Characterization of dynamic patterns of human fetal to neonatal brain asymmetry with deformation-based morphometry

**DOI:** 10.1101/2023.10.30.564508

**Authors:** Céline Steger, Charles Moatti, Kelly Payette, De Silvestro Alexandra, Thi Dao Nguyen, Seline Coraj, Ninib Yakoub, Giancarlo Natalucci, Raimund Kottke, Ruth Tuura, Walter Knirsch, Andras Jakab

## Abstract

Despite established knowledge on the morphological and functional asymmetries in the human brain, the understanding of how brain asymmetry patterns change during late fetal to neonatal life remains incomplete. The goal of this study was to characterize the dynamic patterns of inter-hemispheric brain asymmetry over this critically important developmental stage using longitudinally acquired MRI scans. Super-resolution reconstructed T2-weighted MRI of 20 neurotypically developing participants were used, and for each participant fetal and neonatal MRI was acquired. To quantify brain morphological changes, deformation-based morphometry (DBM) on the longitudinal MRI scans was utilized. Two registration frameworks were evaluated and used in our study: (A) fetal to neonatal image registration and (B) registration through a mid-time template. Developmental changes of cerebral asymmetry were characterized as (A) the inter-hemispheric differences of the Jacobian determinant (JD) of fetal to neonatal morphometry change and the (B) time-dependent change of the JD capturing left-right differences at fetal or neonatal time points. Left-right and fetal-neonatal differences were statistically tested using multivariate linear models, corrected for participants’ age and sex and using threshold-free cluster enhancement. Fetal to neonatal morphometry changes demonstrated asymmetry in the temporal pole and left-right asymmetry differences between fetal and neonatal timepoints revealed temporal changes in the temporal pole, likely to go from right dominant in fetal to a bilateral morphology in neonatal timepoint. Furthermore, the analysis revealed right-dominant subcortical grey matter in neonates and three clusters of increased JD values in the left hemisphere from fetal to neonatal timepoints. While these findings provide evidence that morphological asymmetry gradually emerges during development, discrepancies between registration frameworks require careful considerations when using DBM for longitudinal data of early brain development.

## 1 Introduction

Asymmetry of the brain manifests on various levels, including genetic, morphological, and functional dimensions (Wan et al. 2022). From the neuroscience perspective, studying anatomical asymmetries and their development can reveal insights into the emergence of brain functional specialization, using macro-morphological inter-hemispheric asymmetry as a proxy. The lateralization of sensory-motor functions, behavioral and cognitive processes in the brain is a well-described phenomenon, which is not exclusive to humans. It is widely believed that this lateralization provides evolutionary advantages at the behavioral level, including decreased reaction time and parallel processing in both hemispheres (Güntürkün, Ströckens, and Ocklenburg 2020; Rogers, Zucca, and Vallortigara 2004; Dongen 2006). On a morphological level, post-mortem studies in humans provide evidence for cytoarchitectonic asymmetries (Galaburda, Sanides, and Geschwind 1978) and macroscopic asymmetries such as the ‘Yakovlevian-Torque’ can be observed by pure visual inspection. From the clinical perspective, there is emerging evidence for the association of altered or disrupted hemispheric asymmetry in neurological disorders and neuropsychological conditions (Postema et al. 2019; Sone et al. 2022; Lubben et al. 2021).

Brain asymmetry is likely not a static or linearly emerging property during development; instead, it might involve a complex dynamic process where patterns of left-right asymmetry may appear or diminish due to variations in the expression of developmental genetic programs that control the formation of the characteristic cortical surface patters and brain specialization. Inter-hemispheric asymmetry already develops in the embryonic stage and is induced not only by genetic components but also dependent on environmental factors (Schmitz, Güntürkün, and Ocklenburg 2019; Güntürkün, Ströckens, and Ocklenburg 2020). Post-mortem studies have indicated that right hemispheric structures and a leftward planum temporale (PT) asymmetry have an early origin in fetal brain development (Chi et al., 1977; Wada et al., 1975). While some works found no morphological asymmetries in the fetus using post-mortem analysis (Zhang et al. 2013), asymmetrical cortical folding patterns were described in-vivo as early as the 22^nd^ gestational week (Habas et al. 2012). Several studies described volumetric changes throughout late fetal development for each hemisphere separately to assess their asymmetry (Vasung et al. 2020; Machado-Rivas et al. 2022; Kienast et al. 2021; Cai et al. 2020; Yun et al. 2022). For example, temporal lobe asymmetries are known to change dynamically during the second half of gestation (Kasprian et al. 2011). Asymmetries in the PT and superior temporal sulcus are believed to persist throughout infancy and remain relatively stable until adulthood in healthy human brains (Dubois et al., 2009; Hill et al., 2010).

In addition to the neuroscientific significance of inter-hemispheric asymmetries throughout cerebral development, the final stage of this development holds considerable biological and clinical importance, warranting further investigation in this domain. During the last eight weeks of gestation and early newborn phase, the brain undergoes rapid growth and morphological features, such as secondary and tertiary gyri emerge. In addition, birth takes place, undoubtedly a crucial moment, where a lot of mechanical stress is experienced (Ami et al. 2022) and the entire organism needs to adapt to new conditions and is no longer supported by the mother. Furthermore, this timeframe is of interest, as this is a realistic timeframe to acquire longitudinal data in patients which show altered brain development and risk for brain injuries (Peyvandi et al. 2021) and alternatives to tissue specific volume measurements could provide new insights.

In recent years, advances in fetal and infant MRI have improved access to studying *in* vivo human brain development. However, limited literature exists on longitudinally acquired data and there is a clear limitation of these studies because of their cross-sectional design (that is, each developmental stage is represented by a different individual). Many studies report brain asymmetry, but quantitative MRI data on gestational time-dependent brain morphology changes primarily come from cross-sectional studies, which are constrained by significant inter-individual brain surface variability. Consequently, inferring developmental trends from cross-sectional data can be confounded by this variability and may not accurately represent a within-individual dynamic asymmetry evolution. Furthermore, technical challenges may have contributed to the lack of studies quantifying the dynamic patterns of brain asymmetry over the fetal to neonatal period. Popular neuroscience software and toolboxes are very limited for use with fetal or neonatal MRI data due to different MR image contrasts and not yet developed brain structures, highlighting the urgent need for specialized tools to examine early brain development. While brain volumetry is seen as the gold standard for analyzing brain growth, segmentation of the brain into consistent subregions throughout development remains challenging. Alternative techniques such as deformation-based morphometry (DBM) have been applied before (Ng et al. 2020; Jakab et al. 2019), relying on non-linear image registration. Using DBM, no consistent segmentation of subregions is required to detect local asymmetries of the brain.

The goal of our study was twofold: first, to improve our understanding of the neuroanatomical changes from late fetal to neonatal stage by describing the age-related changes of brain asymmetry based on real longitudinal MRI data acquired at two timepoints. Second, to address outstanding challenges in the deformation-based morphometry analysis of fetal-to-newborn longitudinal development. Specifically, by conducting our analysis using two plausible registration frameworks and evaluating the effects of image post processing options on the image registration accuracy.

## 2 Methods

### 2.1 Participants

Data from twenty healthy participants recruited as participants of two ongoing, prospective studies (BrainDNIU study and BrainCHD study) between 2017 and 2022 were taken for this analysis. Based on patient history, ultrasound (US) and MR scans, all fetuses and newborns were classified as having normal brain development. The female/male ratio was 14/6. All participants were scanned twice: during fetal life and as a newborn. The gestational week (GW) at scan was calculated based on clinical US records. Mean ± standard deviation (SD) age at scan was 32.4 ±1.1 GW for the fetal timepoint and 41.7 ±1.4 GW for neonatal. All babies were born around term at a mean age of 39.7 ±1.2 GW. The captured time interval between the two MRIs was 65 ±12 days for each participant.

Parents gave written informed consent for the participation of the study they were enrolled in as well as consent for data sharing between the studies. BrainCHD and brainDNIU study were approved by the local ethics committee (BASEC IDs: 2019-01993, 2017-00885).

### 2.2 MRI acquisition

Fetal MRI was performed on a 1.5T clinical whole-body MRI scanner (GE Signa Discovery MR450) using a 32-channel cardiac coil or an 8-channel body array coil. The MRI scanner was upgraded to GE Signa Artist during the data collection. Post-upgrade fetal scans were performed with an AIR coil, with the anterior array (blanked coil) having 30 channels and table coil 40 channels. T2-weighted single shot Fast Spin Echo (ssFSE) sequences were acquired in axial, sagittal and coronal orientations, relative to the fetal brain. In-plane resolution was 0.5 mm * 0.5 mm and a slice thickness of 3-5 mm was applied, depending on the covered organ (placenta, whole fetus or brain only). The sequence parameters were the following: TR: 2000-3500 ms, TE: 120 ms (minimum), flip angle: 90. During the scan, the quality of the ssFSE scans was checked by the attending pediatric radiologist and were repeated in case excessive movement or other artifacts were detected. As a rule of thumb, at least two image sequences per orientation were acquired.

In the neonates, all MR scans were performed in natural sleep, without sedation. Neonates received protective earplugs and earmuffs. T2-weighted fast spin-echo (FSE) sequences in all three planes were acquired on a 3.0 T MRI scanner using an 8-channel head coil (GE Signa Discovery MR750). The sequence parameters were the following: TR: 5900 ms, TE: 97 ms, flip angle: 90°. In-plane resolution was 0.35 * 0.35 mm, slice thickness: 2.5 mm with a 0.2 mm slice gap. For the neonatal MRI, only one image sequence per orientation was acquired. During data collection time, the MR scanner was upgraded to GE Signa Premier. Post-upgrade neonatal scans were performed with a 48-channel head coil.

### 2.3 Image pre-processing

The MR images were super-resolution (SR) reconstructed using the slice-to-volume reconstruction toolkit (Kuklisova-Murgasova et al. 2012). Fetal ssFSE images were first checked visually for quality, and at least one axial, sagittal and coronal image was selected for SR reconstruction. Similarly, for the neonatal data, one image from each of the three orientations were used. A reference image was chosen and a mask covering the area of the brain was manually drawn using 3D Slicer Software. Images were pre-processed with N4ITK filter in the 3D Slicer Software (Fedorov et al. 2012) and denoising BTK Toolkit (Rousseau et al. 2013). Both fetal and neonatal images were reconstructed to 0.75 mm isotropic voxel resolution. SR images were reoriented to an age-matched template using a 6 degrees-of-freedom transformation to ensure consistent anatomical orientation. In a final step, brains were skull-stripped using a binary mask based on brain segmentation labels (described below).

As part of the pre-processing for non-linear image registration, we explored whether replacing the black background voxels with bright voxels would affect the performance of the deformable image registration. The aim of this procedure was to reduce the confounding effect of the interface between dark voxels, such as the skull and background air, and the bright voxels of extracerebral fluid spaces, as the latter appear very different between the fetal and neonatal MR images. This step was performed by an in-house developed script with image processing steps that rely on FSL (Jenkinson et al. 2012) and AFNI (Cox 1996). Brain and background classes were identified using 3dkmeans and the background was replaced by the 99^th^ percentile value of the image. See Figure 1 for a fetal and neonatal image example with and without a “bleached” background.

**Figure 1.**
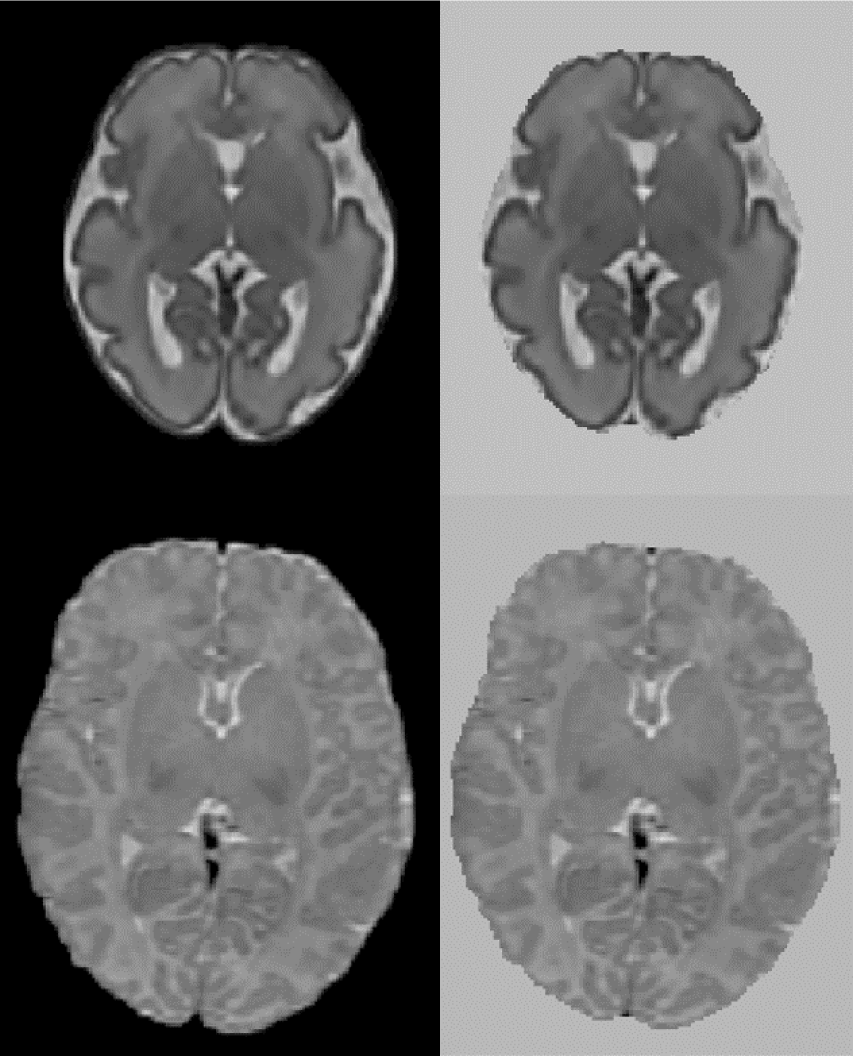
Axial view of fetal and neonatal image of a participant. Top: fetal, bottom: neonatal. Left original image, right image background replaced with bright values.

### 2.4 Characterization of the temporal dynamics of morphological brain asymmetry

To capture time-dependent changes of the inter-hemispheric asymmetry, multiple image registration frameworks are viable choices. Figure 2 gives an overview of the two frameworks used in our study to perform the asymmetry analysis.

**Figure 2.**
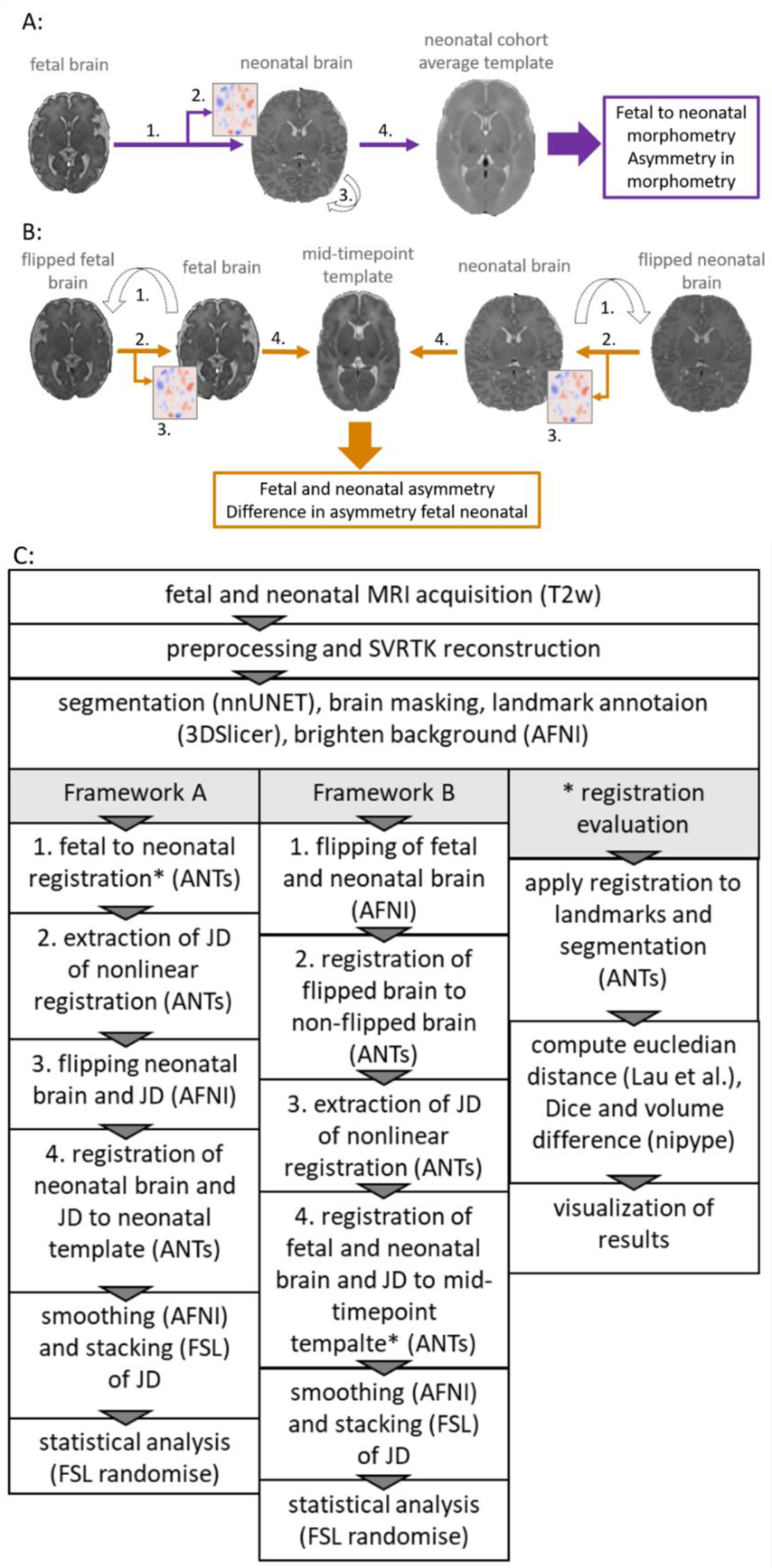
Overview of the two frameworks used to reveal asymmetry patterns and their temporal dynamics. A: framework A (violet), calculating the inter-hemispheric asymmetry of fetal to neonatal regional brain growth maps. B: framework B (orange), calculating fetal and neonatal inter-hemispheric asymmetry at the given timepoint, registering these to a mid-timepoint template and then calculating the differences between the inter-hemispheric asymmetry. C: Flowchart showing steps required to extract JD maps and perform asymmetry analysis for framework A and B and steps required to extract metrics for the registration evaluation.

In framework A (Figure 2 A), we first defined brain growth based on image registrations that map the fetal to the neonatal image within the same participant, directly capturing dynamic morphometric changes from fetal to neonatal timepoints. Similar to Jakab et al. (Jakab et al. 2019), longitudinal MR image pairs were registered for each participant using subsequent linear and non-linear image registrations. A Jacobian determinant (JD) map was calculated from the nonlinear registration component of this step. To statistically test inter-hemispheric asymmetry, as a next step, the JD maps and neonatal MRI were then flipped along the midline axis. The flipped and non-flipped brain and JD pairs were registered to a neonatal template where the asymmetry analysis was performed. Inter-hemispheric difference in morphometric changes were tested as the differences between the flipped and non-flipped JD in the shared template space.

In framework B (Figure 2 B), we first defined static inter-hemispheric asymmetries at the fetal and neonatal time points individually, and then captured their time-dependent changes after registering the asymmetry maps to a mid-age template. For the asymmetry analysis in framework B, fetal and neonatal images were first flipped along the midline axis for each participant. Each flipped image was then registered to the non-flipped image and the JD capturing left-right differences were extracted. Next, the images and JD maps corresponding to the fetal and neonatal timepoints were registered to a mid-age template. Asymmetry analysis was performed by statistically testing for differences between the fetal and neonatal JD maps.

All registration steps were done using the ANTs toolbox (Tustison et al. 2021). Figure 2 C shows an overview of the steps, more details can be found in the Supplementary Material.

### 2.5 Evaluation of registration accuracy for large registration steps

Registration accuracy was tested using two volumetric (image segmentation-based) accuracy metrics and one anatomical landmark-based metric. The rationale for this was to obtain a set of quantitative measurements of accuracy. In the supplementary section we provide an overview of the different registration parameters tried.

#### 2.5.1 Anatomical overlap and volume differences

The fetal and neonatal SR images and the mid-age atlas MRI template were segmented into the following structures: cerebral spinal fluid (CSF), grey matter (GM), white matter (WM), deep grey matter (dGM), brain stem (BS), and cerebellum (Cer). To automate this step, two in-house trained nnUNets (Isensee et al. 2020) were used, corresponding to fetal and neonatal MRI segmentation tasks. The fetal brain segmentation network was trained using the FeTA dataset (Payette et al. 2021), which provides manually annotated ground truth data. For the neonatal data, the ground truth data was generated using the developing human connectome project (dHCP) structural pipeline (Makropoulos et al. 2018). To improve the consistency between the anatomical definition of the segmentations in the fetal and neonatal timepoints, several modifications of the tissue label maps were necessary. We combined the cerebral spinal fluid and ventricle labels. The hippocampus label generated by the neonatal network was assigned to grey matter. All segmentations were visually checked by the first author of this study. The segmented anatomical structures are illustrated in Figure 3.

**Figure 3.**
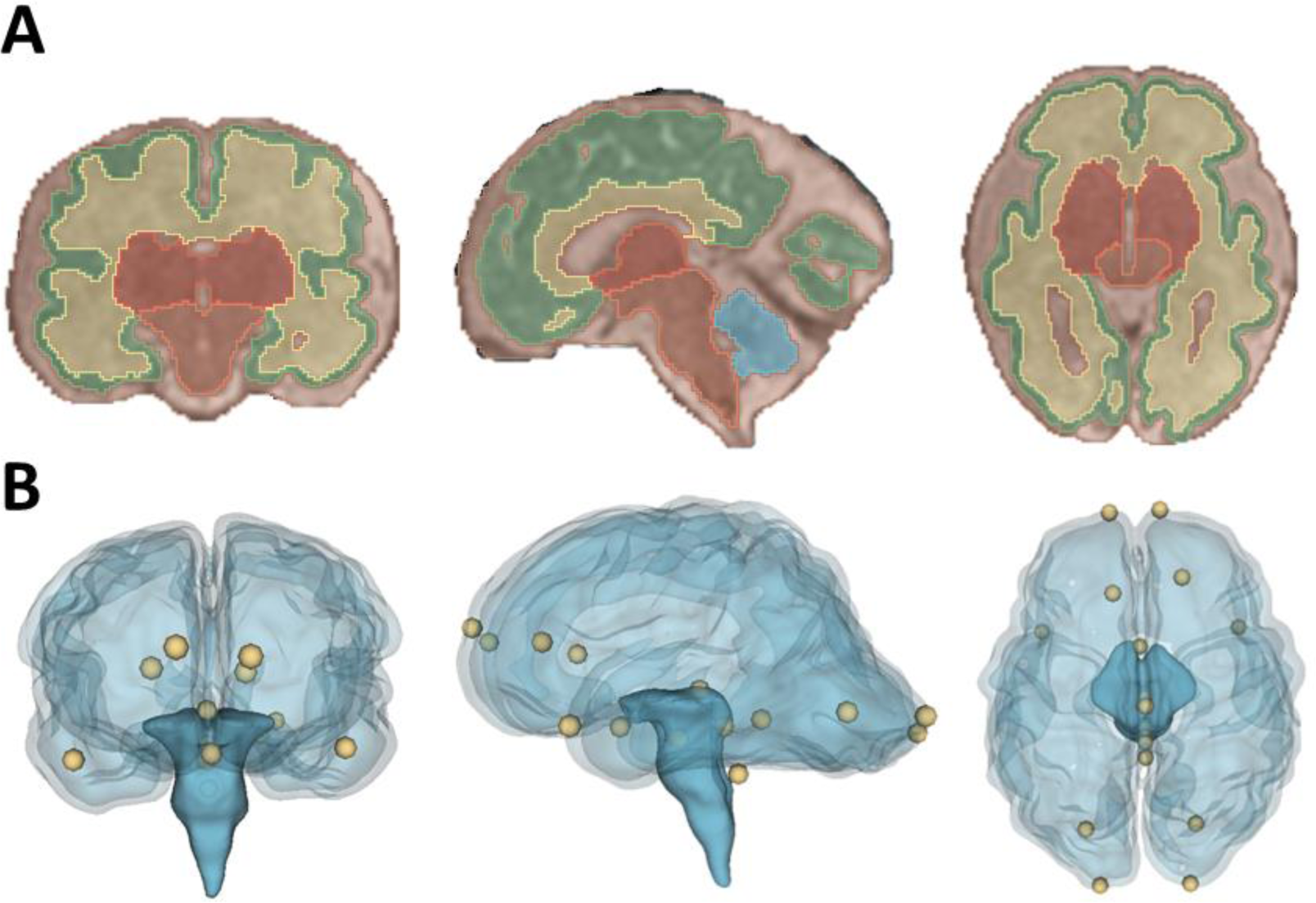
Visualization of the registration evaluation metrics used in our study. A: Coronal, sagittal and axial view of segmentation overlaid on a T2-wieghted SR image of a fetus. B: Coronal, sagittal and axial view of a glass brain rendering showing all sixteen, manually placed landmarks that were used during the registration evaluation.

The two segmentation-based evaluation metrics, Dice overlap coefficient (Dice 1945) and volume difference were evaluated using the co-registered segmentations (framework A: fetal, framework B: fetal and neonatal) and the target segmentation (framework A: participant specific neonatal segmentation, framework B: mid-age template). The Dice coefficient and volume difference were calculated using their implementation in nipype (Gorgolewski et al. 2011). Volume difference is described using the following definition:

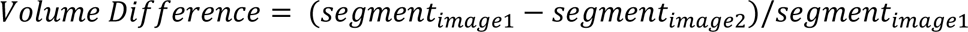

For the evaluation the absolute value of this metric was used.

#### 2.5.2 Anatomical landmark distances

The following landmarks were annotated in each image following Lau (Lau et al. 2019): posterior commissure, infracollicular sulcus, pontomesenphalic junction (PMJ). In addition, the most caudal point of the 4th ventricle and the optic chiasm were annotated and for left and right hemisphere, the rostral end of: hemisphere, temporal lobe and lateral ventricles; and the caudal end of: lateral ventricles and hemisphere were annotated. The landmarks were manually placed in the fetal and neonatal SR images as well as the mid-age template using the Markup module in 3D Slicer, and their coordinates were exported for the registration evaluation. After transforming each landmark with the given registration method, the distance between the ground truth and transformed points were calculated using the Euclidean distance. The anatomical landmarks are illustrated in Figure 3 B.

### 2.6 Statistical analysis of inter-hemispheric brain asymmetry

As defined in section 2.4, both frameworks used JD maps to capture brain morphometry changes or brain asymmetry. Following this logic, the statistical evaluations of brain asymmetry and the time-dependent change of brain asymmetry were performed as follows.

First, JD used for the given analysis were smoothed (using 3dBlurInMask (Cox 1996)) and stacked into a 4D format using fslmerge (Jenkinson et al. 2012). Statistical brain asymmetry analysis was carried out using the randomise tool of FSL. Randomise is an implementation of randomized non-parametric permutation testing (Winkler et al. 2014). Paired sample designs were set up. Threshold-free cluster enhancement (TFCE) (Smith and Nichols 2009) was used to correct for multiple comparisons, using TFCE with 5000 permutations. In framework A the linear regression was performed contrasting two groups: flipped and non-flipped JD maps. Demeaned GW at fetal and neonatal scan were added as covariates. Similarly, in framework B the asymmetry analysis for the fetal and neonatal timepoint was performed. The two groups flipped and non-flipped were contrasted and the demeaned GW at scan and sex were added as covariate. To capture the temporal changes in asymmetry from fetal to neonatal in framework B the (non-flipped) fetal and (non-flipped) neonatal JD were tested for significant differences while including sex and the age variable of both timepoints as covariates.

### 2.7 Visualization and clusters

For visualizing the results of the statistical tests, the anatomical clusters were shown after thresholding them at a value of 0.975, corresponding to p<0.025. FSL Cluster reported the cluster size and generated corresponding labels. Mean, minimum and maximum JD value for each cluster were extracted using the labels. All visualizations were done in fsleyes in the FSL tool, images in the axial view were mirrored to show then in neurological orientation (left hemisphere corresponding to left side in the image). Statistical images were overlaid on the corresponding T2-weighted template images.

## 3 Results

### 3.1 Registration accuracy

In Figure 1 the achieved registration accuracy for the three registrations fetal to neonatal, fetal to mid-template and neonatal to mid-template are visualized. Post-hoc Wilcox test between registration showed significant difference between mean landmarks accuracy of fetal to neonatal and neonatal to template registration (p=0.001). For the mean dice there was a significant difference for all comparisons (fet2neo vs. fetal: p=0.03, fet2neo vs. neonatal: p<0.001, fetal vs. neonatal: p<0.001). Finally, mean volume difference was significantly different for fetal to neonatal registration compared to fetal (p<0.001) and neonatal (p<0.001) to template registration.

**Figure 1a.**
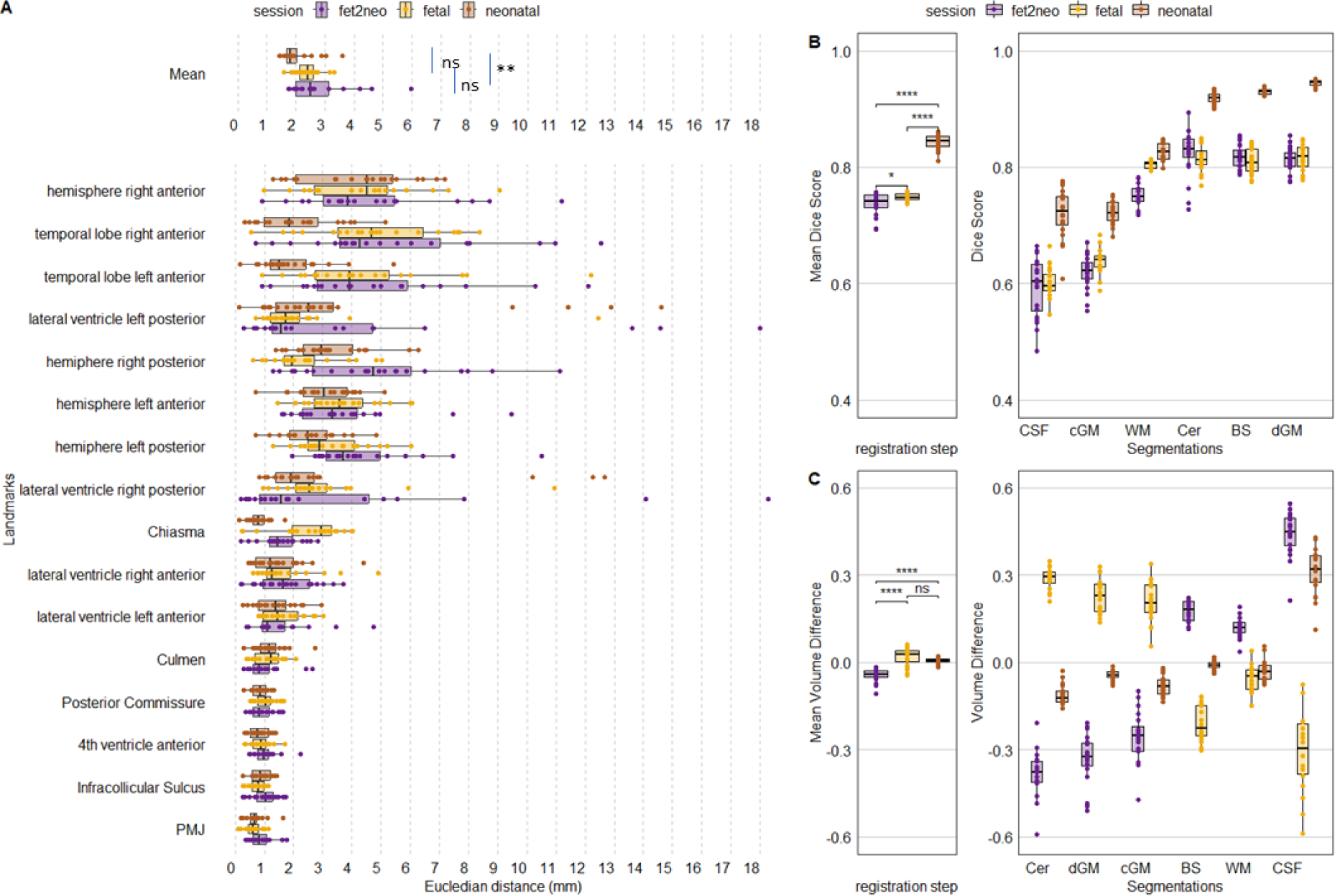
Achieved metric accuracy for each participant for each registration step. A) Landmark accuracy as Euclidean distance in mm. Overall mean landmark distance per registration step (Mean, top row) and individual landmark distance. B) Overall mean Dice score and Dice per segmentation label. C) Overall mean volume difference and volume difference per segmentation label. fet2neo = fetal to neonatal registration, fetal = fetal to template registration, neonatal = neonatal to template registration. PMJ = pontomesenphalic junction.

In addition, a pattern within the individual metrics seems to be visible. Landmarks located around the brainstem seem to have achieved consistently high alignment, whereas other landmarks had larger errors. For the segmentation-based metrics, the Cer, BS and dGM achieved the highest dice score and there was lower accuracy for cGM and CSF. While the mean volume difference was close to zero, the individual labels had larger errors in both negative and positive directions.

Results of a complementary analysis, where we tested different settings for the registration can be found in the Supplementary Materials.

### 3.2 Dynamic patterns of brain morphometry and asymmetry

In all experiments, overall scaling, translation, and rotation from the linear registration component has been factored out since our analyses relied on the deformation component of the fetal to neonatal or mid-age template registration. Therefore, we refer to inter-hemispheric asymmetry as the asymmetry of morphometric change, meaning relative expansion or contraction of a given anatomical structure.

#### 3.2.1 Inter-hemispheric asymmetry of regional fetal to neonatal brain growth maps (framework A)

We first quantified inter-hemispheric differences in regional fetal to neonatal morphometric change using the framework A. A repeated measures linear model (paired flipped vs. non-flipped measurements) revealed two anatomical clusters in which inter-hemispheric differences in the regional fetal to neonatal morphometry change maps were significant (Figure 2). The larger cluster (Table 1, FN 1) was localized to the left temporal pole region extending into cortical gray matter of the temporal lobe as well as the surrounding CSF. The mean values of the JD maps for the flipped and non-flipped hemispheres are visualized within the cluster in the axial view (Figure 2, left panel). This analysis revealed that the regional fetal to neonatal morphometry change map in the left hemisphere (non-flipped) is dominated by positive JD values, referring to regional expansion, however, due to the smoothing applied to the JD maps, it’s not possible to determine whether voxels in the CSF or GM are driving these results.

**Figure 2a.**
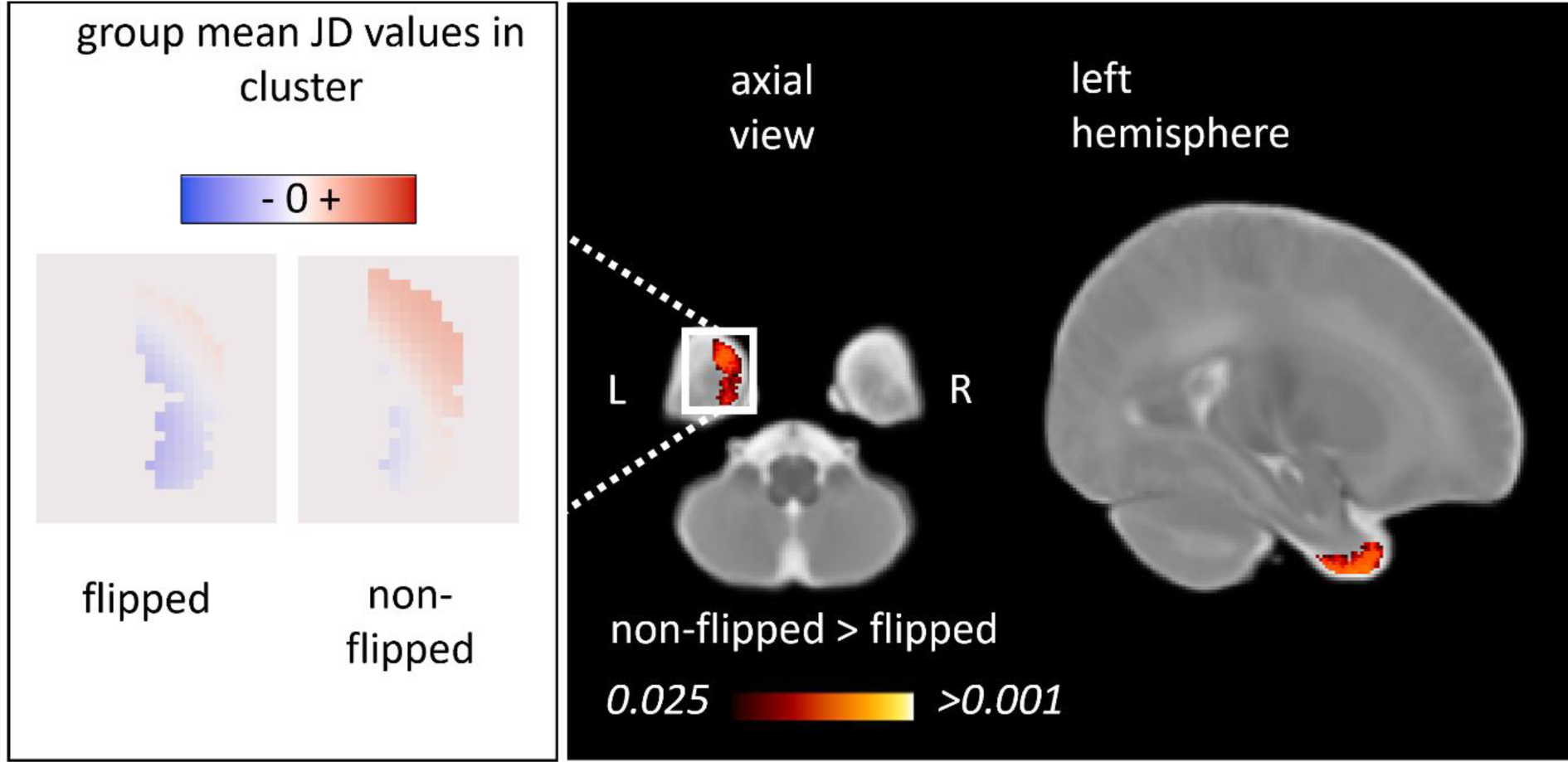
Inter-hemispheric asymmetry of the regional fetal to neonatal morphometry change maps (framework A). We illustrated the anatomical clusters in which flipped and non-flipped morphometry change maps were significantly different, referring to an inter-hemispheric difference in the regional fetal to neonatal morphometry change (TFCE-corrected p<0.025). Left panel: population mean, voxel-level JD values in the left and right hemispheres characterizing regional brain growth (flipped to illustrate anatomical match). Right panel: anatomical clusters overlaid on a neonatal T2-weighted template.

**Table 1.** Anatomical regions demonstrating inter-hemispheric morphometric asymmetry: fetal to neonatal asymmetry analysis (Framework A).

| Cluster | hemisphere | Anatomical location | size (# voxels) | lowest p-value | JD values ipsilateral |  |
| --- | --- | --- | --- | --- | --- | --- |
|  |  |  |  |  | min - max | mean |
| FN1 | Left | Temporal pole | 1462 | 0.006 | (-1.00) – 0.82 | 0.09 |

Clusters of a statistical threshold of TFCE-corrected $p < 0.025$ were included in this table. The clusters represent areas, where the JD values of the non-flipped images were significantly larger compared to the flipped images.

#### 3.2.2 Fetal to neonatal changes of inter-hemispheric asymmetry (framework B)

In framework B, inter-hemispheric asymmetry at each timepoint (fetal or neonatal) was tested after transforming the JD maps to a mid-age template as shown in Figure 6. This approach enabled the quantification of inter-hemispheric asymmetry individually at the fetal and neonatal timepoint as well as the characterization of age-dependent changes. Using a repeated measures linear model (fetal vs. neonatal measurements) regions with larger JD values in neonates were identified. This statistical analysis revealed three anatomical regions where inter-hemispheric asymmetry changed from fetal to neonatal timepoints. Table 2 summarizes the anatomical locations, cluster sizes, minimum p-values as well as JD (fetal and neonatal timepoints) of each cluster. Furthermore, we describe the localization of these clusters and link them to the previously described static asymmetries at the fetal and neonatal age to better characterize the dynamic changes from fetal to neonatal age. Cluster IDs are numbered in descending order of size.

**Figure 3a.**
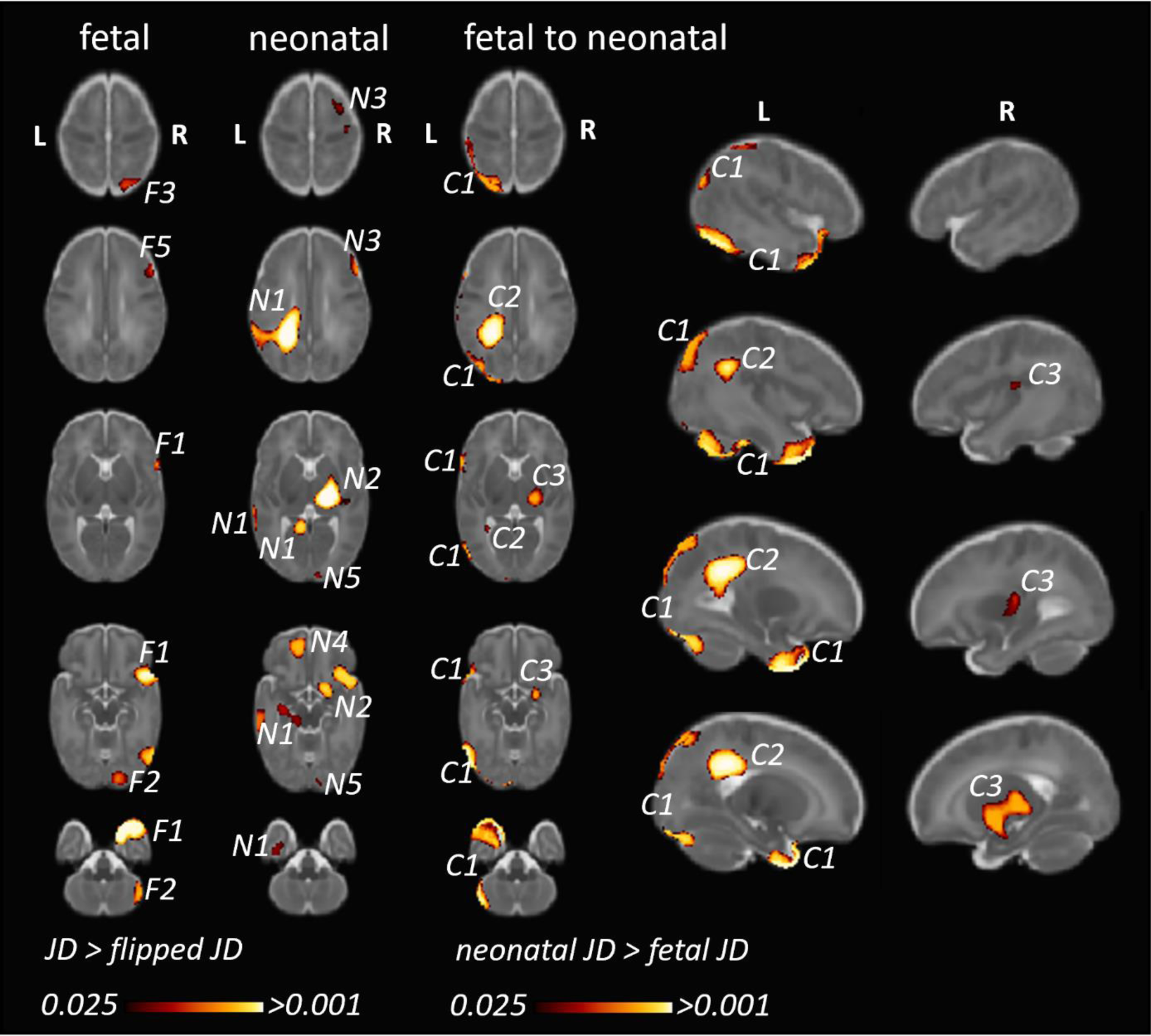
Fetal and neonatal inter-hemispheric asymmetry and dynamic, age-related changes of inter-hemispheric asymmetry (framework B). A: Clusters corresponding to anatomical regions showing (static) inter-hemispheric asymmetry in either the fetal or neonatal time points. B: Clusters corresponding to anatomical regions where inter-hemispheric asymmetry showed differences between the neonatal and fetal timepoints. We only illustrate clusters where neonatal JD was higher than fetal JD, since the opposite statistical test (fetal JD > neonatal) would reveal very similar findings in the homotopic ipsilateral anatomical regions. F1…N: clusters representing regional asymmetry in the fetal images (significant TFCE-corrected clusters where flipped JD>non-flipped JD). N1…N: clusters representing regional asymmetry in the neonatal images (significant TFCE-corrected clusters where flipped JD>non-flipped JD). C1…N: clusters where inter-hemispheric asymmetry changed from fetal to neonatal age.

**Table 2.** Anatomical regions demonstrating inter-hemispheric morphometric asymmetry: fetal and neonatal MRI to template analysis (Framework B).

| C | H | Anatomical location | Corr. cluster | size (# voxels) | lowest p-value | JD values fetal |  | JD values neonatal |  |
| --- | --- | --- | --- | --- | --- | --- | --- | --- | --- |
|  |  |  |  |  |  | min - max | mean | min - max | mean |
| **C1** | left | temporal pole, temporo-occipital cortex, postcentral gyrus, white matter of the parietal cortex | F1, F2, N2 | 27513 | <0.001 | (-1.33) - 0.67 | -0.19 | (-0.71) - 0.46 | -0.02 |
| **C2** | left | centrum semiovale, white matter of the postcentral and angular gyri, left ventricle frontal horn | N1 | 8825 | <0.001 | (-0.66) - 0.60 | -0.07 | (-0.67) - 0.37 | 0.06 |
| **C3** | right | thalamus, basal ganglia, claustrum, internal and external capsules, posterior most aspect of the lateral sulcus | N2 | 5639 | 0.006 | (-0.32) - 0.24 | -0.05 | (-0.15) - 0.29 | 0.05 |

*C=cluster, H = hemisphere; corr. cluster = most likely corresponding cluster*
*The clusters represent areas, where the JD values of the neonatal group were significantly larger compared to the fetal group. Clusters were ordered according to descending size.*

In the left temporal lobe, a large cluster (ID in Table 2 and Figure 6: C1) was localized to the temporal pole, extending superiorly into the temporo-occipital cortex as well as postcentral gyrus in the parietal cortex. Within this cluster, a right-dominant temporal pole was shown at the fetal stage, while no right dominance was found in the neonatal age, therefore the neonatal JD > fetal JD was significant in the left hemisphere for the temporal pole region (Figure 6 C1). In the parietal cortex, extending into the deep white matter of the postcentral gyrus and superior parietal lobule, more leftwards inter-hemispheric asymmetry was demonstrated in the neonates (Figure 6, N1/N4/N7). This marks a trend towards more bilateral morphology of the temporal lobes and temporo-occipital cortex in newborns compared to fetuses (or a left to right shift during fetal to neonatal development), and a trend towards leftwards asymmetry of the deep white matter of the postcentral gyrus and superior temporal lobule (or a right to left shift during fetal to neonatal development).

In the parietal lobe, a cluster was found in the deep cerebral white matter, extending into to the body of the left lateral ventricle and extending superiorly into the parietal white matter of the postcentral and angular gyri (ID in Table 2 and Figure 6: C2). Within the parietal aspect of this cluster, no fetal asymmetry was found (Figure 6), while strong, left dominant neonatal asymmetry was found in the static analysis (N1 in Figure 6). This refers to a trend towards more lateralized morphology in neonatal age in the left hemisphere.

In the deep white matter and subcortical gray matter, a cluster was revealed with its central aspect in the right thalamus, basal ganglia, claustrum, internal and external capsules, extending laterally to the posterior insula / posterior aspect of the central sulcus (ID in Table 2 and Figure 6: C3). In the fetal images, this cluster did not show inter-hemispheric asymmetry (Figure 6), referring to a trend towards more lateralized morphology in neonatal age in the right hemisphere.

While the static asymmetry pattern provided insights into what could have driven the observed dynamic patterns, they also revealed regions that did not correspond to the temporal analysis. In the analysis of the static neonatal asymmetry, cluster N3, located in the right frontal lobe, may correspond to parts of F1 of the fetal asymmetry analysis. The presence of these two clusters could indicate, that asymmetry is present in both timepoints to a similar degree. For F3 located in the occipital lobe and N6 (not visualized) located in the right cerebellum we could not find any correspondence between the time points. As these clusters were only identified in one either fetal or neonatal static asymmetry analysis, but not in the temporal dynamics, their temporal dynamics remain unclear.

#### 3.2.3 Common findings in framework A and framework B

Our working hypothesis was that the two methodological frameworks would yield comparable descriptions of the temporal dynamics of brain asymmetry, this way, cross-validating the results by using two different pipelines. The trend towards more bilateral morphology or a left to right shift of the pole of the left temporal lobe was the only consistent finding across the two image analysis frameworks in our study. However, in both approaches that investigated the temporal dynamics we found anatomical regions that showed different asymmetry patterns in late fetal than newborn age. In framework A the left temporal pole had higher JD values than the right, similarly, in framework B the left temporal pole revealed an asymmetry difference between fetal and neonatal timepoint. When looking at the static asymmetry results, this temporal dynamic seems to be driven by the smaller fetal left temporal pole, which is less asymmetric in the neonatal brains.

## 4 Discussion

Prenatal human brain development is a complex and precisely orchestrated process influenced by genetic programs and environmental factors. The appearance of macroscopic asymmetries between the brain’s hemispheres likely arises from a combination of various developmental events. To gain a deeper understanding of this phenomenon, our study examined the changing patterns of inter-hemispheric brain asymmetry from late fetal development to the neonatal stage. We employed an analytical approach with multiple tasks to optimize the image processing steps, all aimed at answering the fundamental neurobiological question: How does inter-hemispheric asymmetry change in the relatively short period from before birth to after birth? Out of the four areas showing significant differences in inter-hemispheric asymmetry between the two developmental stages, two were predominantly localized to the left temporal lobe and left parietal and postcentral region, one in the right deep gray matter and white matter structures, and the smallest cluster found in the left inferior frontal gyrus. Contrary to our expectation of obtaining concordant results with the two frameworks, only one anatomical region (temporal lobe) survived the TFCE-based correction for multiplicity in two different frameworks of analysis.

### 4.1 The dynamic patterns of inter-hemispheric brain asymmetry over the fetal to neonatal developmental period

We have identified multiple brain regions that exhibit an increasing inter-hemispheric morphometric asymmetry from the fetal to neonatal timepoints. In our analysis, such a region could be detected either as a significant cluster found in both the late fetal and neonatal periods, which gradually became more lateralized as indicated by larger JD values in the ipsilateral hemisphere, or alternatively, a region that is not (significantly) detected in the fetal, but in the neonatal stage.

Specifically, the left-sided deep white matter of the postcentral gyrus, superior parietal lobule, superior temporal gyrus, and planum temporale region present in the dynamic analysis (Figure 6, C2 cluster) was present in both the fetal (Figure 6, F4 cluster) and neonatal brain (Figure 6, N1 cluster). The most concordant finding in our study in terms of across-method reproducibility, cluster size or significance was the inter-hemispheric morphometric asymmetries of the temporal lobe. A cluster in the temporal pole in framework B (C1) was confluent with the postcentral gyrus as well as deep white matter while the temporal pole was the only region to be consistently found in framework A and B.

The parietal lobe is known to display global leftwards dominance, which might be concordant with the leftwards localization of our clusters (C1, C2) in the deep white matter of the superior parietal lobule and angular gyrus (Lehtola et al. 2019). Several studies reported inter-hemispheric asymmetries in the temporal lobe during development, predominantly revealing an emerging left dominance for the planum temporale region and temporal lobe and Sylvian fissure length (Lyttelton et al. 2009; Mallela et al. 2020), but a rightwards dominance in terms of gyral and sulcal development (Chi, Dooling, and Gilles 1977; Rajagopalan et al. 2012, 2011; Kasprian et al. 2011; Habas et al. 2012). The results of multiple studies in infants are consistent with those found in adult brains (Kong et al. 2018), indicating that the structural differences responsible for hemispheric speech and language dominance may be formed during the final trimester of fetal development (Chi, Dooling, and Gilles 1977; Wada and Davis 1977; Gilmore et al. 2007; Hill et al. 2010; Li et al. 2014). The literature regarding the temporal lobe and its development remains ambiguous since an overall rightwards asymmetry for the temporal lobe in newborns has been reported (Lehtola et al. 2019), and there is converging evidence for rightwards superior temporal sulcus asymmetry (Hill et al. 2010; Leroy et al. 2015; Specht and Wigglesworth 2018). Our study, however, used longitudinal data, therefore removing some of the ambiguity of inter-individual variability that may hinder studies that concluded developmental changes of asymmetry based on cross-sectional data. A similar study using a longitudinal design and a versatile, surface-based analysis in preterm infants replicated well-known asymmetries in the Sylvian fissure and a more significant opercularization in the left hemisphere, in contrast to the previously documented early sulcation development in the right hemisphere (De Vareilles et al. 2023).

We identified a cluster with right sided (increasing) neonatal morphometric lateralization located in the subcortical gray matter, internal and external capsules, extending ventrally into the hippocampus or the temporal horn of the right lateral ventricle. Our finding is concordant with one of others, suggesting that morphological features of subcortical grey matter in neonates are right dominant (Dean et al. 2018; Ratnarajah et al. 2013). However, in the largest performed study of this kind using the data of the ENIGMA consortium, the thalamus and basal ganglia were overall left dominant, while the caudate nucleus, hippocampus and amygdala were right dominant (Guadalupe et al. 2017). While the cluster C3/N2 reached ventrally to the hippocampus, it is too early to conclude whether this is a finding concordant with the data from the ENIGMA study, since we also revealed a strong rightwards dominance in the neonatal brain (N2) in a region that encompasses the lateral aspects of the thalamus. It is highly likely that asymmetries in the thalamus reflect asymmetry of the white matter pathways they relay, and therefore might not be consistently left or right dominant for all parts of the thalamus and basal ganglia.

The smallest cluster showing dynamically changing inter-hemispheric asymmetry was localized to the left frontal lobe, around the region of the left inferior frontal gyrus. Data from adults in the ENIGMA consortium confirmed rightwards asymmetry of the surface area of the inferior frontal gyrus, however, there is a scarcity of developmental studies confirming this observation. While this is the least significant finding in our analysis, it is possible that it marks an emerging leftwards dominance of the inferior frontal gyrus.

There are several ambiguities in our findings. For instance, there are regions that were statistically significantly lateralized in the late fetal images but not in the neonatal images (such as F3, F5, F6, F7, see Table 3 and Figure 6) or regions that were only detected in the neonatal images (such as N4, N6 and N7, see Table 3 and Figure 6). There are several reasons for these changes. First, technical aspects such as the presence of clusters that are smaller than the registration inaccuracy or noise differed between fetal and neonatal images. Secondly, there is the possibility that very subtle changes did not survive the statistical correction for multiple comparisons.

**Table 3.** Anatomical regions demonstrating inter-hemispheric morphometric asymmetry, static asymmetry in the fetal and neonatal timepoints.

| C | H | Anatomical location | corr. clusters | size (# voxels) | lowest p-value | JD values ipsilateral |  |
| --- | --- | --- | --- | --- | --- | --- | --- |
|  |  |  |  |  |  | min – max | mean |
| Fetal inter-hemispheric morphometric asymmetry |  |  |  |  |  |  |  |
| F1 | right | superior frontal gyrus, temporal pole | N2, C1 | 13191 | <0.001 | -0.31 – 1.03 | 0.15 |
| F2 | right | occipital pole, cerebellar hemisphere | C1 | 6654 | 0.003 | -0.52 – 0.74 | 0.13 |
| F3 | right | occipital cortex, cuneus | C1 | 1995 | 0.013 | -0.64 – 0.62 | 0.13 |
| F4 | right | Superior frontal gyrus | N3 | 1515 | 0.013 | -0.27 – 0.57 | 0.11 |
| F5 | right | Superior frontal gyrus | N3 | 273 | 0.02 | -0.26 – 0.65 | 0.8 |
| F6 | left | occipital cortex, primary visual | N1 | 178 | 0.023 | -0.13 - 0.28 | 0.09 |
| Neonatal inter-hemispheric morphometric asymmetry |  |  |  |  |  |  |  |
| N1 | left | white matter, postcentral gyrus, superior parietal lobule, superior temporal gyrus / PT region | C2 | 22230 | <0.001 | -0.61 – 0.40 | 0.05 |
| N2 | right | thalamus, basal ganglia, claustrum, internal and external capsules, posteriormost aspect of the lateral sulcus | F1, C1 | 18602 | <0.001 | -0.54 – 0.49 | 0.06 |
| N3 | right | superior frontal gyrus | F5, F4 | 5606 | 0.007 | -0.22 – 0.60 | 0.04 |
| N4 | left | olfactory cortex, white matter of the superior frontal gyrus |  | 4230 | 0.005 | -0.18 – 0.26 | 0.04 |
| N5 | right | occipital pole |  | 815 | 0.017 | (-0.66) - 0.49 | 0.12 |
| N6 | right | cerebellum |  | 506 | 0.013 | -0.07 – 0.29 | 0.07 |
| N7 | left | Deep white matter of the frontal lobe |  | 407 | 0.023 | -0.13 - 0.22 | 0.22<sup>0.05</sup> |

*C=cluster, H = hemisphere with the larger JD value; corr. cluster = most likely corresponding cluster* *The clusters represent areas, where the JD values on the ipsilateral side (non-flipped image) were larger than on the contralateral side (flipped image). Clusters were ordered according to descending size.*

### 4.2 Selection of registration methods for DBM in longitudinal fetal to neonatal cohorts

For DBM, accurate registration is critical for obtaining reliable results. A lower degree of anatomical correspondence would require larger spatial smoothing, which is a crucial confounding factor in DBM, VBM and related studies (Scarpazza et al. 2015). Evaluation of registration accuracy was crucially important for our study as the size and morphology of the brain changes significantly from late fetal to newborn stage. Different frameworks may offer varying levels of precision in the registration process. In the next section, we provide a critical review and highlight the potential limitations of our methodology to select the best registration method for morphometry.

While visual inspection of the results is often advised, it still can be challenging to assess the best registration strategy for the given problem. Here, we used a set of landmarks and segmentation-based metrics to quantify the registration accuracy. Segmentation based metrics, similarly to volumetric studies, rely on consistent segmentation across the two timepoints. Therefore, we used tissue labels, which are easier to segment consistently across fetal and neonatal brains but allow for less local evaluation of the accuracy than finer segmentation labels. The distance between two landmarks on the other hand is a single point and therefore describes the accuracy on a very local level. However, it is challenging to establish a set of suitable landmarks, as they need to be visible in the images in both timepoints.

After nonlinear registration, the achieved accuracy was moderate (Mean Dice of 0.75, 0.85 and 0.74 and mean landmark distance of 2.4, 2.0 and 2.8 mm, respectively for fetal to template, neonatal to template and fetal to neonatal registration). These results were achieved by increasing the number of iterations, reducing the radius for the cross-correlation to two and replacing the background of the fetal images with a bright value (See Supplementary section Registration testing).

When using segmentation and landmark based metrics, certain errors have to be expected. Oishi et al. (Oishi et al. 2011) performed registration accuracy tests for their atlas based anatomical segmentation of neonatal brains. The average Dice measure was 0.82, which was similar to the achieved accuracy using manual segmentation. However, the segmented structures were anatomically detailed which makes a direct comparison to our metrics not possible. Similarly, for the landmark-based metric, Lau et al. (Lau et al. 2019) developed a landmark based framework to evaluate the anatomical correspondence between to brain images in adults. They reported a landmark localization error of less than 1 mm for landmarks such as PMJ, culmen and infracollicular sulcus and a mean error after nonlinear registration of 0.68, 2.7 and 0.93. For our worst performing registration setting, fetal to neonatal, we could report registration for the PMJ of 1.07 after linear and 0.82 after nonlinear registration. While some of the landmarks were inspired by the paper of Lau et al. (Lau et al. 2019), we also identified our own landmarks that were further away from the brain stem which might have suffered from greater inaccuracy during the annotation. The motivation behind this was to capture the registration accuracy in the entire brain and not just core structures.

Our results imply that static asymmetry in the neonatal stage may be more easily and reliably detected. This is likely due to the larger size, better image quality, increased development and maturity of the neonatal brain, which makes it easier to identify morphometric features even with less precise registration.

### 4.3 Methodological limitations

Our findings on inter-hemispheric brain asymmetry are limited by the following methodological factors. In framework A, the large registration from the fetal to neonatal timepoints is challenging, however the JD which describes morphometric changes within each participant already captures the longitudinal aspect of the data. It describes the change between the two timepoints and simply by flipping the JD maps it is possible to perform longitudinal asymmetry testing. In framework B, the asymmetry at each timepoint is captured by registering the flipped to the non-flipped brain. By registration to a shared template the asymmetry then can be tested for differences. The registration to a shared template could be seen as less challenging as the timeframe and therefore difference between participant and template might be smaller. However, it is important to consider that the accuracy of the neonatal to template registration is higher than that for the fetal to template registration. Another inherent aspect of this framework is that the generation of JD maps based on flipped fetal to non-flipped fetal and flipped and non-flipped neonatal brains might produce subtle systematic differences due to the difference in the images.

In our study, we utilized image registrations in DBM, to infer inter-hemispheric asymmetry. However, this approach has some limitations with regard to the interpretability of our findings. We referred to our statistically significant “clusters” as anatomical regions where the morphometric features of the brain differ between homotopic regions of the two hemispheres. This approach is somewhat restrictive, as it relies on information contained within the JD maps to infer morphology. It is worth noting that the JD maps are based on image-based registration (and not surface or landmark-based), which aims to map visual features between a moving and target image.

Consequently, the expansion or contraction of a brain region is only detected if there are features that can be matched and which expand or contract. In many brain regions, this is not the case during development. For instance, for the cortex, we primarily detected shrinkage (negative JD values) as the general tendency from fetal to neonatal age, which may be attributed to the increasing convolutional pattern (gyrification) with age after factoring out the majority of global brain scaling that occurred during the linear registration step. Furthermore, our non-linear registration methods are not likely to capture emerging structures such as two gyri appearing instead of one, or structures rotating, etc. As a result, while we employ a robust statistical approach and a dual-framework analysis, we cannot definitively conclude whether the asymmetries are driven by (asymmetric) isotropic growth, folding, rotation, or other types of morphometric differences between the two hemispheres.

A possible solution to these problems would be to perform landmark, sulcal, surface-based analyses (Lefevre et al. 2016; Lebenberg et al. 2018; Ahmad et al. 2019; Bozek et al. 2018; Oishi, Chang, and Huang 2019), however, there the fidelity and topological correctness of meshes and segmentation accuracy in fetal brain remains a critical challenge (de Dumast et al. 2022; Payette et al. 2022). A newer approach might be to learn registrations by deep learning (Wei et al. 2020), however, it is not yet clear how efficient these networks are for fetal and neonatal development. Alternatively, a future approach could be to re-visit each region individually in each participant and provide an observer based manual annotations (such as in (Kasprian et al. 2011)) to reveal concrete morphometric changes.

The present study is subject to limitations due to the relatively small sample size (n=20), limiting the statistical power of our analyses. Nevertheless, this challenge of achieving substantial sample sizes is present in many studies using fetal or neonatal MRI of healthy pregnancies. To substantiate our findings, we suggest that validation of our results using comparable datasets or a database comprising preterm newborns, where clinically indicated longitudinal MRI scans are available.

### 4.4 Conclusion

In conclusion, our study complements existing neuroscientific studies on inter-hemispheric neuroanatomical asymmetry by providing evidence that morphological asymmetry is not static during late fetal development, and dynamic processes take place during late fetal development. Importantly, the neuroscientific value of our work is to characterize the dynamic patterns of asymmetry within-subject, therefore reducing some of the confounding effects of inter-individual variability that is present in cross-sectional studies on human brain development. Our results underscore the importance of the perinatal period for brain development, encompassing significant milestones such as tertiary gyri emergence and maturation, myelination, and cortical adaptation in response to external stimuli. The study also implies that a suitable image registration is critical when using DBM and that results might vary when using different frameworks, which must be considered when interpreting results.

## Acknowledgments

The authors would like to thank all families that participated in the BrainCHD and BrainDNIU study and clinical staff involved in the data collection. We especially thank the radiology and neonatology department of the University Children’s Hospital Zurich.

## 5 Data Availability Statement

Original imaging dataset are not readily available due to restrictions in consent. Non-participant specific data supporting the conclusions of this article will be made available by the authors, without undue reservation. Code is available on GitHub: https://github.com/CST333/fet2neoasymmetry

## References

Ahmad, Sahar, Zhengwang Wu, Gang Li, Li Wang, Weili Lin, Pew-Thian Yap, and Dinggang Shen. 2019. “Surface-Constrained Volumetric Registration for the Early Developing Brain.” Medical Image Analysis 58 (101540): 101540.

Ami, Olivier, Jean-Christophe Maran, Dominique Musset, Claude Dubray, Gerard Mage, and Louis Boyer. 2022. “Using Magnetic Resonance Imaging During Childbirth to Demonstrate Fetal Head Moldability and Brain Compression: Prospective Cohort Study.” JMIR Formative Research 6 (11): e27421.

Bozek, Jelena, Antonios Makropoulos, Andreas Schuh, Sean Fitzgibbon, Robert Wright, Matthew F. Glasser, Timothy S. Coalson, et al. 2018. “Construction of a Neonatal Cortical Surface Atlas Using Multimodal Surface Matching in the Developing Human Connectome Project.” NeuroImage 179 (October): 11–29.

Cai, Shulei, Guofu Zhang, He Zhang, and Jing Wang. 2020. “Normative Linear and Volumetric Biometric Measurements of Fetal Brain Development in Magnetic Resonance Imaging.” Child’s Nervous System: ChNS: Official Journal of the International Society for Pediatric Neurosurgery 36 (12): 2997–3005.

Chi, J. G., E. C. Dooling, and F. H. Gilles. 1977. “Left-Right Asymmetries of the Temporal Speech Areas of the Human Fetus.” Archives of Neurology 34 (6): 346–48.

Cox, R. W. 1996. “AFNI: Software for Analysis and Visualization of Functional Magnetic Resonance Neuroimages.” Computers and Biomedical Research, an International Journal 29 (3): 162–73.

De Vareilles, Heloise, Denis Riviere, Marco Pascucci, Zhong-Yi Sun, Clara Fischer, Francois Leroy, Maria-Luisa Tataranno, Manon J. Benders, Jessica Dubois, and Jean-Francois Mangin. 2023. “Exploring the Emergence of Morphological Asymmetries around the Brain’s Sylvian Fissure: A Longitudinal Study of Shape Variability in Preterm Infants.” Cerebral Cortex, January. 10.1093/cercor/bhac533.

Dean, Douglas C., III, E. M. Planalp, W. Wooten, C. K. Schmidt, S. R. Kecskemeti, C. Frye, N. L. Schmidt, H. H. Goldsmith, A. L. Alexander, and R. J. Davidson. 2018. “Correction to: Investigation of Brain Structure in the 1-Month Infant.” Brain Structure & Function 223 (6): 3007–9.

Dice, Lee R. 1945. “Measures of the Amount of Ecologic Association between Species.” Ecology 26 (3): 297–302.

Dongen, S. V. 2006. “Fluctuating Asymmetry and Developmental Instability in Evolutionary Biology: Past, Present and Future.” Journal of Evolutionary Biology 19 (6): 1727–43.

Dumast, Priscille de, Hamza Kebiri, Vincent Dunet, Meriam Koob, and Meritxell Bach Cuadra. 2022. “Multi-Dimensional Topological Loss for Cortical Plate Segmentation in Fetal Brain MRI.” ArXiv [Eess.IV]. arXiv. http://arxiv.org/abs/2208.07566.

Fedorov, Andriy, Reinhard Beichel, Jayashree Kalpathy-Cramer, Julien Finet, Jean-Christophe Fillion-Robin, Sonia Pujol, Christian Bauer, et al. 2012. “3D Slicer as an Image Computing Platform for the Quantitative Imaging Network.” Magnetic Resonance Imaging 30 (9): 1323–41.

Galaburda, A. M., F. Sanides, and N. Geschwind. 1978. “Human Brain. Cytoarchitectonic Left-Right Asymmetries in the Temporal Speech Region.” Archives of Neurology 35 (12): 812–17.

Gilmore, John H., Weili Lin, Marcel W. Prastawa, Christopher B. Looney, Y. Sampath, K. Vetsa, Rebecca C. Knickmeyer, Dianne D. Evans, et al. 2007. “Regional Gray Matter Growth, Sexual Dimorphism, and Cerebral Asymmetry in the Neonatal Brain.” The Journal of Neuroscience: The Official Journal of the Society for Neuroscience 27 (6): 1255–60.

Gorgolewski, Krzysztof, Christopher D. Burns, Cindee Madison, Dav Clark, Yaroslav O. Halchenko, Michael L. Waskom, and Satrajit S. Ghosh. 2011. “Nipype: A Flexible, Lightweight and Extensible Neuroimaging Data Processing Framework in Python.” Frontiers in Neuroinformatics. 10.3389/fninf.2011.00013.

Guadalupe, Tulio, Samuel R. Mathias, Theo G. M. vanErp, Christopher D. Whelan, Marcel P. Zwiers, Yoshinari Abe, Lucija Abramovic, et al. 2017. “Human Subcortical Brain Asymmetries in 15,847 People Worldwide Reveal Effects of Age and Sex.” Brain Imaging and Behavior 11 (5): 1497–1514.

GUnturkun, Onur, Felix Ströckens, and Sebastian Ocklenburg. 2020. “Brain Lateralization: A Comparative Perspective.” Physiological Reviews 100 (3): 1019–63.

Habas, Piotr A., Julia A. Scott, Ahmad Roosta, Vidya Rajagopalan, Kio Kim, Francois Rousseau, A. James Barkovich, Orit A. Glenn, and Colin Studholme. 2012. “Early Folding Patterns and Asymmetries of the Normal Human Brain Detected from in Utero MRI.” Cerebral Cortex 22 (1): 13–25.

Hill, Jason, Donna Dierker, Jeffrey Neil, Terrie Inder, Andrew Knutsen, John Harwell, Timothy Coalson, and David Van Essen. 2010. “A Surface-Based Analysis of Hemispheric Asymmetries and Folding of Cerebral Cortex in Term-Born Human Infants.” The Journal of Neuroscience: The Official Journal of the Society for Neuroscience 30 (6): 2268–76.

Isensee, Fabian, Paul F. Jaeger, Simon A. A. Kohl, Jens Petersen, and Klaus H. Maier-Hein. 2020. “NnU-Net: A Self-Configuring Method for Deep Learning-Based Biomedical Image Segmentation.” Nature Methods 18 (2): 203–11.

Jakab, Andras, Eliane Meuwly, Maria Feldmann, Michael von Rhein, Raimund Kottke, Ruth O’Gorman Tuura, Beatrice Latal, Walter Knirsch, and Research Group Heart and Brain. 2019. “Left Temporal Plane Growth Predicts Language Development in Newborns with Congenital Heart Disease.” Brain: A Journal of Neurology 142 (5): 1270–81.

Jenkinson, Mark, Christian F. Beckmann, Timothy E. J. Behrens, Mark W. Woolrich, and Stephen M. Smith. 2012. “FSL.” NeuroImage 62 (2): 782–90.

Kasprian, Gregor, Georg Langs, Peter C. Brugger, Mario Bittner, Michael Weber, Mavilde Arantes, and Daniela Prayer. 2011. “The Prenatal Origin of Hemispheric Asymmetry: An in Utero Neuroimaging Study.” Cerebral Cortex (New York, N.Y.: 1991) 21 (5): 1076–83.

Kienast, Patric, Ernst Schwartz, Mariana C. Diogo, Gerlinde M. Gruber, Peter C. Brugger, Herbert Kiss, Barbara Ulm, et al. 2021. “The Prenatal Origins of Human Brain Asymmetry: Lessons Learned from a Cohort of Fetuses with Body Lateralization Defects.” Cerebral Cortex 31 (8): 3713–22.

Kong, Xiang-Zhen, Samuel R. Mathias, Tulio Guadalupe, David C. Glahn, Barbara Franke, Fabrice Crivello, Nathalie Tzourio-Mazoyer, et al. 2018. “Mapping Cortical Brain Asymmetry in 17,141 Healthy Individuals Worldwide via the ENIGMA Consortium.” Proceedings of the National Academy of Sciences of the United States of America 115 (22): E5154–63.

Kuklisova-Murgasova, Maria, Gerardine Quaghebeur, Mary A. Rutherford, Joseph V. Hajnal, and Julia A. Schnabel. 2012. “Reconstruction of Fetal Brain MRI with Intensity Matching and Complete Outlier Removal.” Medical Image Analysis 16 (8): 1550–64.

Lau, Jonathan C., Andrew G. Parrent, John Demarco, Geetika Gupta, Jason Kai, Olivia W. Stanley, Tristan Kuehn, et al. 2019. “A Framework for Evaluating Correspondence between Brain Images Using Anatomical Fiducials.” Human Brain Mapping 40 (14): 4163–79.

Lebenberg, J., M. Labit, G. Auzias, H. Mohlberg, C. Fischer, D. Riviere, E. Duchesnay, et al. 2018. “A Framework Based on Sulcal Constraints to Align Preterm, Infant and Adult Human Brain Images Acquired in Vivo and Post Mortem.” Brain Structure & Function 223 (9): 4153–68.

Lefevre, Julien, David Germanaud, Jessica Dubois, Francois Rousseau, Ines de Macedo Santos, Hugo Angleys, Jean-Francois Mangin, Petra S. Huppi, Nadine Girard, and Francois De Guio. 2016. “Are Developmental Trajectories of Cortical Folding Comparable Between Cross-Sectional Datasets of Fetuses and Preterm Newborns?” Cerebral Cortex 26 (7): 3023–35.

Lehtola, S. J., J. J. Tuulari, L. Karlsson, R. Parkkola, H. Merisaari, J. Saunavaara, T. Lahdesmaki, N. M. Scheinin, and H. Karlsson. 2019. “Associations of Age and Sex with Brain Volumes and Asymmetry in 2-5-Week-Old Infants.” Brain Structure & Function 224 (1): 501–13.

Leroy, Francois, Qing Cai, Stephanie L. Bogart, Jessica Dubois, Olivier Coulon, Karla Monzalvo, Clara Fischer, et al. 2015. “New Human-Specific Brain Landmark: The Depth Asymmetry of Superior Temporal Sulcus.” Proceedings of the National Academy of Sciences of the United States of America 112 (4): 1208–13.

Li, Gang, Jingxin Nie, Li Wang, Feng Shi, Amanda E. Lyall, Weili Lin, John H. Gilmore, and Dinggang Shen. 2014. “Mapping Longitudinal Hemispheric Structural Asymmetries of the Human Cerebral Cortex from Birth to 2 Years of Age.” Cerebral Cortex (New York, N.Y.: 1991) 24 (5): 1289–1300.

Lubben, Noah, Elizabeth Ensink, Gerhard A. Coetzee, and Viviane Labrie. 2021. “The Enigma and Implications of Brain Hemispheric Asymmetry in Neurodegenerative Diseases.” Brain Communications 3 (3): fcab211.

Lyttelton, Oliver C., Sherif Karama, Yasser Ad-Dab’bagh, Robert J. Zatorre, Felix Carbonell, Keith Worsley, and Alan C. Evans. 2009. “Positional and Surface Area Asymmetry of the Human Cerebral Cortex.” NeuroImage 46 (4): 895–903.

Machado-Rivas, Fedel, Jasmine Gandhi, Jungwhan John Choi, Clemente Velasco-Annis, Onur Afacan, Simon K. Warfield, Ali Gholipour, and Camilo Jaimes. 2022. “Normal Growth, Sexual Dimorphism, and Lateral Asymmetries at Fetal Brain MRI.” Radiology 303 (1): 162–70.

Makropoulos, Antonios, Emma C. Robinson, Andreas Schuh, Robert Wright, Sean Fitzgibbon, Jelena Bozek, Serena J. Counsell, et al. 2018. “The Developing Human Connectome Project: A Minimal Processing Pipeline for Neonatal Cortical Surface Reconstruction.” NeuroImage 173 (June): 88–112.

Mallela, Arka N., Hansen Deng, Alyssa K. Brisbin, Alan Bush, and Ezequiel Goldschmidt. 2020. “Sylvian Fissure Development Is Linked to Differential Genetic Expression in the Pre-Folded Brain.” Scientific Reports 10 (1): 14489.

Ng, Isabel H. X., Alexandra F. Bonthrone, Christopher J. Kelly, Lucilio Cordero-Grande, Emer J. Hughes, Anthony N. Price, Jana Hutter, et al. 2020. “Investigating Altered Brain Development in Infants with Congenital Heart Disease Using Tensor-Based Morphometry.” Scientific Reports 10 (1): 14909.

Oishi, Kenichi, Linda Chang, and Hao Huang. 2019. “Baby Brain Atlases.” NeuroImage 185 (January): 865–80.

Oishi, Kenichi, Susumu Mori, Pamela K. Donohue, Thomas Ernst, Lynn Anderson, Steven Buchthal, Andreia Faria, et al. 2011. “Multi-Contrast Human Neonatal Brain Atlas: Application to Normal Neonate Development Analysis.” NeuroImage 56 (1): 8–20.

Payette, Kelly, Priscille de Dumast, Hamza Kebiri, Ivan Ezhov, Johannes C. Paetzold, Suprosanna Shit, Asim Iqbal, et al. 2021. “An Automatic Multi-Tissue Human Fetal Brain Segmentation Benchmark Using the Fetal Tissue Annotation Dataset.” Scientific Data 8 (1): 167.

Payette, Kelly, Hongwei Li, Priscille de Dumast, Roxane Licandro, Hui Ji, Md Mahfuzur Rahman Siddiquee, Daguang Xu, et al. 2022. “Fetal Brain Tissue Annotation and Segmentation Challenge Results.” ArXiv [Eess.IV]. arXiv. http://arxiv.org/abs/2204.09573.

Peyvandi, Shabnam, Jessie Mei Lim, Davide Marini, Duan Xu, V. Mohan Reddy, A. James Barkovich, Steven Miller, Patrick McQuillen, and Mike Seed. 2021. “Fetal Brain Growth and Risk of Postnatal White Matter Injury in Critical Congenital Heart Disease.” The Journal of Thoracic and Cardiovascular Surgery 162 (3): 1007–1014.e1.

Postema, Merel C., Daan van Rooij, Evdokia Anagnostou, Celso Arango, Guillaume Auzias, Marlene Behrmann, Geraldo Busatto Filho, et al. 2019. “Altered Structural Brain Asymmetry in Autism Spectrum Disorder in a Study of 54 Datasets.” Nature Communications 10 (1): 4958.

Rajagopalan, Vidya, Julia Scott, Piotr A. Habas, Kio Kim, James Corbett-Detig, Francois Rousseau, A. James Barkovich, Orit A. Glenn, and Colin Studholme. 2011. “Local Tissue Growth Patterns Underlying Normal Fetal Human Brain Gyrification Quantified in Utero.” The Journal of Neuroscience: The Official Journal of the Society for Neuroscience 31 (8): 2878–87.

Rajagopalan, Vidya, Julia Scott, Piotr A. Habas, Kio Kim, Francois Rousseau, Orit A. Glenn, A. James Barkovich, and Colin Studholme. 2012. “Mapping Directionality Specific Volume Changes Using Tensor Based Morphometry: An Application to the Study of Gyrogenesis and Lateralization of the Human Fetal Brain.” NeuroImage 63 (2): 947–58.

Ratnarajah, Nagulan, Anne Rifkin-Graboi, Marielle V. Fortier, Yap Seng Chong, Kenneth Kwek, Seang-Mei Saw, Keith M. Godfrey, Peter D. Gluckman, Michael J. Meaney, and Anqi Qiu. 2013. “Structural Connectivity Asymmetry in the Neonatal Brain.” NeuroImage 75 (July): 187–94.

Rogers, Lesley J., Paolo Zucca, and Giorgio Vallortigara. 2004. “Advantages of Having a Lateralized Brain.” Proceedings. Biological Sciences 271 Suppl 6 (suppl_6): S420–2.

Rousseau, Francois, Estanislao Oubel, Julien Pontabry, Marc Schweitzer, Colin Studholme, Meriam Koob, and Jean-Louis Dietemann. 2013. “BTK: An Open-Source Toolkit for Fetal Brain MR Image Processing.” Computer Methods and Programs in Biomedicine 109 (1): 65–73.

Scarpazza, Cristina, Stefania Tognin, Silvia Frisciata, Giuseppe Sartori, and Andrea Mechelli. 2015. “False Positive Rates in Voxel-Based Morphometry Studies of the Human Brain: Should We Be Worried?” Neuroscience and Biobehavioral Reviews 52 (May): 49–55.

Schmitz, Judith, Onur Gunturkun, and Sebastian Ocklenburg. 2019. “Building an Asymmetrical Brain: The Molecular Perspective.” Frontiers in Psychology 10 (April): 982.

Smith, Stephen M., and Thomas E. Nichols. 2009. “Threshold-Free Cluster Enhancement: Addressing Problems of Smoothing, Threshold Dependence and Localisation in Cluster Inference.” NeuroImage 44 (1): 83–98.

Sone, Mari, Daisuke Koshiyama, Yinghan Zhu, Norihide Maikusa, Naohiro Okada, Osamu Abe, Hidenori Yamasue, Kiyoto Kasai, and Shinsuke Koike. 2022. “Structural Brain Abnormalities in Schizophrenia Patients with a History and Presence of Auditory Verbal Hallucination.” Translational Psychiatry 12 (1): 511.

Specht, Karsten, and Philip Wigglesworth. 2018. “The Functional and Structural Asymmetries of the Superior Temporal Sulcus.” Scandinavian Journal of Psychology 59 (1): 74–82.

Tustison, Nicholas J., Philip A. Cook, Andrew J. Holbrook, Hans J. Johnson, John Muschelli, Gabriel A. Devenyi, Jeffrey T. Duda, et al. 2021. “The ANTsX Ecosystem for Quantitative Biological and Medical Imaging.” Scientific Reports 11 (1): 9068.

Vasung, Lana, Caitlin K. Rollins, Hyuk Jin Yun, Clemente Velasco-Annis, Jennings Zhang, Konrad Wagstyl, Alan Evans, et al. 2020. “Quantitative In Vivo MRI Assessment of Structural Asymmetries and Sexual Dimorphism of Transient Fetal Compartments in the Human Brain.” Cerebral Cortex 30 (3): 1752–67.

Wada, J. A., and A. E. Davis. 1977. “Fundamental Nature of Human Infant’s Brain Asymmetry.” The Canadian Journal of Neurological Sciences. Le Journal Canadien Des Sciences Neurologiques 4 (3): 203–7.

Wan, Bin, $eyma Bayrak, Ting Xu, H. Lina Schaare, Richard A. I. Bethlehem, Boris C. Bernhardt, and Sofie L. Valk. 2022. “Heritability and Cross-Species Comparisons of Human Cortical Functional Organization Asymmetry.” ELife 11 (July): e77215.

Wei, Dongming, Sahar Ahmad, Yunzhi Huang, Lei Ma, Zhengwang Wu, Gang Li, Li Wang, Qian Wang, Pew-Thian Yap, and Dinggang Shen. 2020. “An Auto-Context Deformable Registration Network for Infant Brain MRI.” ArXiv [Cs.CV]. arXiv. http://arxiv.org/abs/2005.09230.

Winkler, Anderson M., Gerard R. Ridgway, Matthew A. Webster, Stephen M. Smith, and Thomas E. Nichols. 2014. “Permutation Inference for the General Linear Model.” NeuroImage 92 (May): 381–97.

Yun, Hyuk Jin, Hyun Ju Lee, Joo Young Lee, Tomo Tarui, Caitlin K. Rollins, Cynthia M. Ortinau, Henry A. Feldman, P. Ellen Grant, and Kiho Im. 2022. “Quantification of Sulcal Emergence Timing and Its Variability in Early Fetal Life: Hemispheric Asymmetry and Sex Difference.” NeuroImage, September, 119629.

Zhang, Z., Z. Hou, X. Lin, G. Teng, H. Meng, F. Zang, F. Fang, and S. Liu. 2013. “Development of the Fetal Cerebral Cortex in the Second Trimester: Assessment with 7T Postmortem MR Imaging.” AJNR. American Journal of Neuroradiology 34 (7): 1462–67.

