## Supplemental Material for "Characterization of dynamic patterns of human fetal to neonatal brain asymmetry with deformation-based morphometry"

### Supplementary Material

#### 1 Complementary Analysis: Registration Settings Testing

##### 1.1 Detailed Image registration benchmark

As described in section 2.4, two frameworks were used to capture the dynamic changes of inter-hemispheric brain asymmetry. These required three different types of image registrations: direct fetal to neonatal registration, fetal to mid-template and neonatal to mid-template registration. Each of these consists of a linear and a subsequent non-linear step to ensure accurate anatomical correspondence between the structures. As a next step in our analysis, we evaluated the accuracy of a popular image registration tool with various, commonly used settings.

###### 1.1.1 Linear registration

For the linear registration, we used the linear registration algorithm implemented in the FSL software (flirt) (Jenkinson et al. 2002) and the ANTs software (Tustison et al. 2021). All linear registrations resulted in affine transformations with 12-degrees of freedom. To find a suitable linear registration for our study, the transformation settings in ANTs python implementation 'antspy' as well as settings based on recommendations for the terminal use of the ANTs toolbox were tested. All registrations were run separately with and without masking of the cost function. For the masking, we created two different types of binary masks, one based on the image segmentations directly and an eroded version of this brain mask (generated using fslmaths). We tested the results of an eroded masking strategy as this minimizes the effect of the most outer image intensities which still could contain darker voxels after skull stripping and may create artificial boundaries to the image and affect the alignment and scaling of the linear registration. Detailed information on the various configurations of the linear registrations used can be found in the **Supplementary Table 1**.

###### 1.1.2 Deformable image registration

For the deformable image registration we explored two non-linear registration methods implemented in the ANTs software (Tustison et al. 2021): the Symmetric Normalization (SyN) transformation model (Avants et al. 2008) and the time-varying diffeomorphism with mean square metric. Since there is no recommendation on the optimal configuration for the large deformation that our study might require, we evaluated different combinations of pre-processing options, optimization settings and cost metrics. In this series of tests, we only considered settings that used the binary mask generated from the segmentation as this was also the mask used in the later steps of the asymmetry analysis.

Detailed information on the various settings of the deformable image registrations used can be found in the **Supplementary Table 2.**

##### 1.1.3 Evaluation of registration accuracy

Registration accuracy was tested using two volumetric (image segmentation-based) accuracy metrics and one anatomical landmark-based metric. The rationale for this was to obtain a set of quantitative measurements of accuracy that then can be ranked based on a set of defined rules to find the most suitable registration settings for the entire dataset.

###### 1.1.3.1 Registration evaluation ranking

To rank the different registration algorithms and settings, we used the ChallengeR tool (Wiesenfarth et al. 2021). This R package was originally designed to rank the performance of algorithms in e.g., image segmentation challenges. Similarly, different registration settings were ranked on their performance of a task, e.g., the distance of a specific landmark to its reference. For each evaluated registration step and metric, a ranking was performed using the ranking scheme aggregate using function "mean", then rank. Next, for each registration step, the results of each metric ranking were summed, and an overall winner was defined. To take into consideration that the volume difference and Dice coefficient both reflect volumetric measures, we assigned a weight of 0.5 to them, so that the landmark-based metrics (distance measure) contributed equally to the summed ranking score:

$$summed\ ranking\ score = 0.5 * (consensus\ rank\ dice + consensus\ rank\ volume\ difference) + consensus\ rank\ Landmarks$$

**Supplementary Table 1: Linear registration settings**

| **Naming** | **call** |
|---|---|
| **Terminal/bash CC** | antsRegistration --dimensionality 3 --float 0 \ <br> --output [${warp},${image}] \ <br> --interpolation Linear \ <br> --winsorize-image-intensities [0.005,0.995] \ <br> --use-histogram-matching 1 \ <br> --initial-moving-transform [${template},${subj},1] \ |

| | |
|---|---|
| | --transform Rigid[0.1] \ <br><br> --metric CC[${template},${subj},1,2,Regular,0.25] \ <br><br> --convergence [1000x500x250x100,1e-6,10] \ <br><br> --shrink-factors 8x4x2x1 \ <br><br> --smoothing-sigmas 3x2x1x0 \ <br><br> --transform Affine[0.1] \ <br><br> --metric CC[${template},${subj},1,2,Regular,0.25] \ <br><br> --convergence [1000x500x250x100,1e-6,10] \ <br><br> --shrink-factors 8x4x2x1 \ <br><br> --smoothing-sigmas 3x2x1x0 \ <br><br> -x [${maskfix},${maskmoving}] --verbose 1 <br><br> (see https://github.com/ANTsX/ANTs/wiki/Anatomy-of-an-antsRegistration-call) |
| **Terminal/bash MI** | antsRegistration --dimensionality 3 --float 0 \ <br><br> --output [${warp},${image}] \ <br><br> --interpolation Linear \ <br><br> --winsorize-image-intensities [0.005,0.995] \ <br><br> --use-histogram-matching 1 \ |

| | |
|---|---|
| | --initial-moving-transform [${template},${subj},1] \<br>--transform Rigid[0.1] \<br>--metric MI[${template},${subj},1,32,Regular,0.25] \<br>--convergence [1000x500x250x100,1e-6,10] \<br>--shrink-factors 8x4x2x1 \<br>--smoothing-sigmas 3x2x1x0 \<br>--transform Affine[0.1] \<br>--metric MI[${template},${subj},1,32,Regular,0.25] \<br>--convergence [1000x500x250x100,1e-6,10] \<br>--shrink-factors 8x4x2x1 \<br>--smoothing-sigmas 3x2x1x0 \<br>-x [${maskfix},${maskmoving}] --verbose 1<br>(see https://github.com/ANTsX/ANTs/wiki/Anatomy-of-an-antsRegistration-call) |
| **antspy linear: Affine, TRSAA, Similarity** | #(for transform in ["Affine", "TRSAA", "Similarity"]:)<br>ants.registration(fixed = fixed, moving = moving, mask=mask_temp,<br> type_of_transform= transform ) |
| **antsAI** | antsAI -d 3 \ |

| | |
|---|---|
| | -t Rigid[0.1] \ <br><br> -m GC[${fixed}, ${moving}] \ <br><br> -t Affine[0.1] \ <br><br> -m GC[${fixed}, ${moving}] \ <br><br> -m Mattes[${fixed}, ${moving}, 32, None, 0,3] \ <br><br> -x [${fixedImageMask}, ${movingImageMask}] \ <br><br> -o ${linear_registration} -v 1; |
| **FSL flirt** | flirt -in ${moving} -ref ${fixed} -omat ${linear_registration} -out ${outputfile}; \ <br><br> ./c3d-0.8.2-Linux-x86_64/bin/c3d_affine_tool -ref ${fixed} -src ${moving} ${linear_registration} -fsl2ras -oitk ${linear_registration::-4}_flirt2ants.txt' |

**Supplementary Table 2: Nonlinear registration settings**

| type_of_transform | syn_metric | syn_sampling | reg_iterations (reg_iter, registration iterations) |
|---|---|---|---|
| ‘SyNOnly’ | MI | 32 | (40, 20, 0) (antspy default)<br>(100, 70, 50, 20) |
| ‘SyNOnly’ | CC | 2, 4, 6 | (40, 20, 0) (antspy default)<br>(100, 70, 50, 20) |
| 'TVMSQ' | | | (40, 20, 0) (antspy default)<br>(100, 70, 50, 20) |
| 'TVMSQC' | | | (40, 20, 0) (antspy default)<br>(100, 70, 50, 20) |

**RESUTLS**

In **Supplementary Table 3** the top ranked and chosen linear registration strategies for the asymmetry analysis framework are shown. To linearly align the brains to their target, a TRSAA registration was ranked best for all three registration steps, however with a different masking strategy for each. Overall, the neonatal to template registration achieved better mean metric results suggesting a better linear alignment. **Supplementary Table 4** provides an overview of the highest ranked and therefore chosen registration approach for each of the three registration steps. After initial tests demonstrated that using no mask in the nonlinear step led to registration failure, we continued by only considering approaches that used the “masked” option as this was the area from where the JD is computed. For the neonatal to template registration step two settings ranked equally good, SyN algorithm with increased iterations, cross-correlation as metric and radius 2 and radius 4. Therefore radius 2, the same radius as in the fetal to template registration was chosen. The rankings for the parameter combinations are shown in **Supplementary Tables 5-10**.

**Supplementary Table 3: Linear registration.** For each registration step the winner of the ranking approach was chosen**.**

| **Registration step** | **Linear Registration setting** | **Metric** | **Consensus Rank per Metric** | ***Metric median and IQR*** | **Metric mean and sd** |
|---|---|---|---|---|---|
| fetal to template | TRSAA<br>eroded masked | dice | 1 | 0.671 0.031 | 0.67 0.018 |
| | | volume difference | 2 | -0.042 0.126 | 0.076 0.076 |
| | | landmarks distance | 5 | 3.28 0.56 | 3.239 0.572 |
| neonatal to template | TRSAA<br>masked | dice | 4 | 0.649 0.066 | 0.65 0.039 |
| | | volume difference | 1 | 0.024 0.105 | 0.068 0.068 |
| | | landmarks distance | 1 | 2.979 0.833 | 3.003 0.702 |
| fetal to neonatal | TRSAA<br>nomask | dice | 2 | 0.622 0.038 | 0.627 0.026 |
| | | volume difference | 2 | -0.227 0.069 | -0.222 0.051 |
| | | landmarks distance | 1 | 3.372 1.264 | 3.593 1.046 |

**Supplementary Table 4: Nonlinear registration.** For each registration step the winner of the ranking approach was chosen. In the registration step neonatal to template two settings ranked a tie, SyN algorithm using CC as metric and increased iterations with radius 2 and 4. As in the other registration steps radius 2 performed better, radius 2 was chosen.

| Registration step | Nonlinear Registration setting | Metric | Consensus Rank per Metric | Metric median and IQR | Metric mean and sd |
|---|---|---|---|---|---|
| fetal to template | SyN<br>CC radius 2<br>iterations (100, 70, 50, 20)<br>bleached background | dice | 2 | 0.749 0.01 | 0.748 0.007 |
| | | volume difference | 1 | 0.029 0.037 | 0.019 0.031 |
| | | landmarks distance | 3 | 2.412 0.49 | 2.395 0.424 |
| neonatal to template | SyN<br>CC radius 2*<br>iterations (100, 70, 50, 20) | dice | 1 | 0.846 0.017 | 0.844 0.013 |
| | | volume difference | 1 | 0.007 0.01 | 0.006 0.008 |
| | | landmarks distance | 2 | 1.823 0.342 | 2.018 0.573 |
| fetal to neonatal | SyN<br>CC radius 2<br>iterations (100, 70, 50, 20)<br>bleached background | dice | 1 | 0.742 0.021 | 0.737 0.019 |
| | | volume difference | 2 | -0.043 0.023 | -0.045 0.022 |
| | | landmarks distance | 1 | 2.505 1.129 | 2.796 1.097 |

**Supplementary Table 5: Ranking linear registration settings fetal to template**

| Registration setting tested | meanRanks | consensusrank | metric | summed_consesnsus_ranking | final_ranking |
|---|---|---|---|---|---|
| TRSAA_eroded_masked | 1.833333 | 1 | dice | 6.5 | 1 |
| TRSAA_eroded_masked | 7.5 | 2 | voldiff | 6.5 | 1 |
| TRSAA_eroded_masked | 4.75 | 5 | landmarks | 6.5 | 1 |
| TRSAA_masked | 3.5 | 2 | dice | 7 | 2 |
| TRSAA_masked | 8.166667 | 8 | voldiff | 7 | 2 |
| TRSAA_masked | 3.5 | 2 | landmarks | 7 | 2 |
| Affine_eroded_masked | 3.5 | 2 | dice | 7 | 2 |
| Affine_eroded_masked | 7.666667 | 6 | voldiff | 7 | 2 |
| Affine_eroded_masked | 4.1875 | 3 | landmarks | 7 | 2 |
| TRSAA_nomask | 4 | 4 | dice | 7 | 2 |
| TRSAA_nomask | 7.5 | 2 | voldiff | 7 | 2 |

| | | | | | |
|---|---|---|---|---|---|
| TRSAA_nomask | 4.375 | 4 | landmarks | 7 | 2 |
| Affine_masked | 6 | 6 | dice | 9 | 5 |
| Affine_masked | 9.166667 | 10 | voldiff | 9 | 5 |
| Affine_masked | 3.125 | 1 | landmarks | 9 | 5 |
| Affine_nomask | 5 | 5 | dice | 9.5 | 6 |
| Affine_nomask | 7.5 | 2 | voldiff | 9.5 | 6 |
| Affine_nomask | 5.0625 | 6 | landmarks | 9.5 | 6 |
| Similarity_eroded_masked | 6.166667 | 7 | dice | 11 | 7 |
| Similarity_eroded_masked | 7.333333 | 1 | voldiff | 11 | 7 |
| Similarity_eroded_masked | 6.625 | 7 | landmarks | 11 | 7 |
| Similarity_masked | 7.833333 | 8 | dice | 12 | 8 |
| Similarity_masked | 7.5 | 2 | voldiff | 12 | 8 |
| Similarity_masked | 6.625 | 7 | landmarks | 12 | 8 |
| Similarity_nomask | 8.666667 | 9 | dice | 16.5 | 9 |

| | | | | | |
|---|---|---|---|---|---|
| Similarity_nomask | 7.666667 | 6 | voldiff | 16.5 | 9 |
| Similarity_nomask | 7.5625 | 9 | landmarks | 16.5 | 9 |
| flirt_NA | 9.166667 | 10 | dice | 21.5 | 10 |
| flirt_NA | 8.333333 | 9 | voldiff | 21.5 | 10 |
| flirt_na | 11.8125 | 12 | landmarks | 21.5 | 10 |
| terminal_CC_nomask | 11 | 11 | dice | 21.5 | 10 |
| terminal_CC_nomask | 9.666667 | 12 | voldiff | 21.5 | 10 |
| terminal_CC_nomask | 11 | 10 | landmarks | 21.5 | 10 |
| terminal_CC_masked | 11.333333 | 12 | dice | 23.5 | 12 |
| terminal_CC_masked | 10.5 | 13 | voldiff | 23.5 | 12 |
| terminal_CC_masked | 11.0625 | 11 | landmarks | 23.5 | 12 |
| terminal_MI_nomask | 14.666667 | 14 | dice | 26.5 | 13 |
| terminal_MI_nomask | 9.5 | 11 | voldiff | 26.5 | 13 |
| terminal_MI_nomask | 14.6875 | 14 | landmarks | 26.5 | 13 |

| | | | | | |
|---|---|---|---|---|---|
| terminal_CC_eroded_masked | 13.333333 | 13 | dice | 28 | 14 |
| terminal_CC_eroded_masked | 11.666667 | 17 | voldiff | 28 | 14 |
| terminal_CC_eroded_masked | 12.3125 | 13 | landmarks | 28 | 14 |
| terminal_MI_masked | 14.666667 | 14 | dice | 29 | 15 |
| terminal_MI_masked | 11.333333 | 16 | voldiff | 29 | 15 |
| terminal_MI_masked | 14.6875 | 14 | landmarks | 29 | 15 |
| antsAI_masked | 15.833333 | 16 | dice | 31 | 16 |
| antsAI_masked | 11 | 14 | voldiff | 31 | 16 |
| antsAI_masked | 14.75 | 16 | landmarks | 31 | 16 |
| terminal_MI_eroded_masked | 16.5 | 17 | dice | 32.5 | 17 |
| terminal_MI_eroded_masked | 11 | 14 | voldiff | 32.5 | 17 |
| terminal_MI_eroded_masked | 16.875 | 17 | landmarks | 32.5 | 17 |

**Supplementary Table 6: Ranking linear registration settings neonatal to template**

| algorithm | meanRanks | consensusrank | metric | summed_consesnsus_ranking | final_ranking |
|---|---|---|---|---|---|
| TRSAA_masked | 5.5 | 4 | dice | 3.5 | 1 |
| TRSAA_masked | 5.666667 | 1 | voldiff | 3.5 | 1 |
| TRSAA_masked | 5.3125 | 1 | landmarks | 3.5 | 1 |
| Affine_masked | 5.5 | 4 | dice | 6.5 | 2 |
| Affine_masked | 6.666667 | 5 | voldiff | 6.5 | 2 |
| Affine_masked | 5.375 | 2 | landmarks | 6.5 | 2 |
| Affine_eroded_masked | 6.666667 | 8 | dice | 8 | 3 |
| Affine_eroded_masked | 6 | 2 | voldiff | 8 | 3 |
| Affine_eroded_masked | 5.4375 | 3 | landmarks | 8 | 3 |
| terminal_MI_nomask | 4.833333 | 1 | dice | 11 | 4 |
| terminal_MI_nomask | 8 | 9 | voldiff | 11 | 4 |
| terminal_MI_nomask | 7 | 6 | landmarks | 11 | 4 |
| terminal_MI_masked | 5.166667 | 3 | dice | 11.5 | 5 |

| | | | | | |
|---|---|---|---|---|---|
| terminal_MI_masked | 8.666667 | 10 | voldiff | 11.5 | 5 |
| terminal_MI_masked | 6.625 | 5 | landmarks | 11.5 | 5 |
| TRSAA_eroded_masked | 7.333333 | 9 | dice | 11.5 | 5 |
| TRSAA_eroded_masked | 6.833333 | 6 | voldiff | 11.5 | 5 |
| TRSAA_eroded_masked | 6.375 | 4 | landmarks | 11.5 | 5 |
| terminal_CC_masked | 5 | 2 | dice | 12 | 7 |
| terminal_CC_masked | 7.166667 | 8 | voldiff | 12 | 7 |
| terminal_CC_masked | 8.0625 | 7 | landmarks | 12 | 7 |
| terminal_CC_nomask | 5.833333 | 6 | dice | 12.5 | 8 |
| terminal_CC_nomask | 6.333333 | 3 | voldiff | 12.5 | 8 |
| terminal_CC_nomask | 8.125 | 8 | landmarks | 12.5 | 8 |
| terminal_CC_eroded_masked | 6.333333 | 7 | dice | 19 | 9 |
| terminal_CC_eroded_masked | 9.416667 | 11 | voldiff | 19 | 9 |
| terminal_CC_eroded_masked | 9.125 | 10 | landmarks | 19 | 9 |

| | | | | | |
|---|---|---|---|---|---|
| terminal_MI_eroded_masked | 7.833333 | 10 | dice | 21.5 | 10 |
| terminal_MI_eroded_masked | 12.666667 | 15 | voldiff | 21.5 | 10 |
| terminal_MI_eroded_masked | 9.0625 | 9 | landmarks | 21.5 | 10 |
| Similarity_eroded_masked | 11.666667 | 11 | dice | 23 | 11 |
| Similarity_eroded_masked | 6.333333 | 3 | voldiff | 23 | 11 |
| Similarity_eroded_masked | 13.5625 | 16 | landmarks | 23 | 11 |
| Affine_nomask | 11.833333 | 12 | dice | 23.5 | 12 |
| Affine_nomask | 11.333333 | 13 | voldiff | 23.5 | 12 |
| Affine_nomask | 9.9375 | 11 | landmarks | 23.5 | 12 |
| Similarity_masked | 11.833333 | 12 | dice | 24.5 | 13 |
| Similarity_masked | 7 | 7 | voldiff | 24.5 | 13 |
| Similarity_masked | 12.125 | 15 | landmarks | 24.5 | 13 |
| TRSAA_nomask | 11.833333 | 12 | dice | 25 | 14 |
| TRSAA_nomask | 12 | 14 | voldiff | 25 | 14 |

| | | | | | |
|---|---|---|---|---|---|
| TRSAA_nomask | 10 | 12 | landmarks | 25 | 14 |
| flirt_NA | 13.333333 | 15 | dice | 26.5 | 15 |
| flirt_NA | 11.25 | 12 | voldiff | 26.5 | 15 |
| flirt_na | 11.125 | 13 | landmarks | 26.5 | 15 |
| Similarity_nomask | 15.5 | 16 | dice | 30 | 16 |
| Similarity_nomask | 13.5 | 16 | voldiff | 30 | 16 |
| Similarity_nomask | 11.375 | 14 | landmarks | 30 | 16 |
| antsAI_masked | 17 | 17 | dice | 34 | 17 |
| antsAI_masked | 14.166667 | 17 | voldiff | 34 | 17 |
| antsAI_masked | 14.375 | 17 | landmarks | 34 | 17 |

**Supplementary Table 7: Ranking linear registration settings fetal to neonatal**

| algorithm | meanRanks | consensusrank | metric | summed_consesnsus_ranking | final_ranking |
|---|---|---|---|---|---|
| TRSAA_nomask | 2.833 | 2 | dice | 3 | 1 |
| TRSAA_nomask | 3 | 1 | landmarks | 3 | 1 |

| | | | | | |
|---|---|---|---|---|---|
| TRSAA_nomask | 5.833 | 2 | voldiff | 3 | 1 |
| TRSAA_masked | 3.5 | 4 | dice | 7 | 4 |
| TRSAA_masked | 3.062 | 2 | landmarks | 7 | 4 |
| TRSAA_masked | 6 | 4 | voldiff | 7 | 4 |
| TRSAA_eroded_masked | 4.312 | 5 | landmarks | 8.5 | 5 |
| TRSAA_erroded_masked | 1.5 | 1 | dice | 8.5 | 5 |
| TRSAA_erroded_masked | 6.833 | 6 | voldiff | 8.5 | 5 |
| flirt_NA | 3.167 | 3 | dice | 6.5 | 3 |
| flirt_na | 4.062 | 4 | landmarks | 6.5 | 3 |
| flirt_NA | 5.833 | 2 | voldiff | 6.5 | 3 |
| Affine_nomask | 4 | 5 | dice | 6 | 2 |
| Affine_nomask | 3.688 | 3 | landmarks | 6 | 2 |
| Affine_nomask | 5.667 | 1 | voldiff | 6 | 2 |
| Similarity_nomask | 6.833 | 7 | dice | 12 | 6 |

| | | | | | |
|---|---|---|---|---|---|
| Similarity_nomask | 5.438 | 6 | landmarks | 12 | 6 |
| Similarity_nomask | 6.5 | 5 | voldiff | 12 | 6 |
| Affine_masked | 6.333 | 6 | dice | 13.5 | 7 |
| Affine_masked | 6.5 | 7 | landmarks | 13.5 | 7 |
| Affine_masked | 7 | 7 | voldiff | 13.5 | 7 |
| Similarity_masked | 9 | 9 | dice | 16.5 | 8 |
| Similarity_masked | 8.75 | 8 | landmarks | 16.5 | 8 |
| Similarity_masked | 7.833 | 8 | voldiff | 16.5 | 8 |
| Affine_eroded_masked | 9.188 | 9 | landmarks | 18 | 9 |
| Affine_erroded_masked | 7.833 | 8 | dice | 18 | 9 |
| Affine_erroded_masked | 8.167 | 10 | voldiff | 18 | 9 |
| Similarity_eroded_masked | 10.812 | 11 | landmarks | 20 | 10 |
| Similarity_erroded_masked | 10.5 | 10 | dice | 20 | 10 |
| Similarity_erroded_masked | 7.833 | 8 | voldiff | 20 | 10 |

| | | | | | |
|---|---|---|---|---|---|
| terminal_CC_nomask | 11.5 | 11 | dice | 21 | 11 |
| terminal_CC_nomask | 10.625 | 10 | landmarks | 21 | 11 |
| terminal_CC_nomask | 11 | 11 | voldiff | 21 | 11 |
| terminal_CC_eroded_masked | 11.812 | 13 | landmarks | 26.5 | 12 |
| terminal_CC_erroded_masked | 12.833 | 12 | dice | 26.5 | 12 |
| terminal_CC_erroded_masked | 12.333 | 15 | voldiff | 26.5 | 12 |
| terminal_CC_masked | 13.333 | 13 | dice | 26.5 | 12 |
| terminal_CC_masked | 11.438 | 12 | landmarks | 26.5 | 12 |
| terminal_CC_masked | 13.667 | 16 | voldiff | 26.5 | 12 |
| terminal_MI_eroded_masked | 14.625 | 15 | landmarks | 28 | 14 |
| terminal_MI_erroded_masked | 13.333 | 13 | dice | 28 | 14 |
| terminal_MI_erroded_masked | 11.167 | 13 | voldiff | 28 | 14 |
| terminal_MI_nomask | 14.833 | 15 | dice | 30.5 | 15 |
| terminal_MI_nomask | 15.375 | 16 | landmarks | 30.5 | 15 |

| | | | | | |
|---|---|---|---|---|---|
| terminal_MI_nomask | 12 | 14 | voldiff | 30.5 | 15 |
| antsAI_masked | 16.5 | 17 | dice | 31 | 17 |
| antsAI_masked | 14 | 14 | landmarks | 31 | 17 |
| antsAI_masked | 14.333 | 17 | voldiff | 31 | 17 |
| terminal_MI_masked | 15.167 | 16 | dice | 30.5 | 15 |
| terminal_MI_masked | 16.312 | 17 | landmarks | 30.5 | 15 |
| terminal_MI_masked | 11 | 11 | voldiff | 30.5 | 15 |

**Supplementary Table 8: Ranking nonlinear registration settings fetal to template**

| Registration setting tested | meanRanks | consensusrank | metric | summed_consesnsus_ranking | final_ranking |
|---|---|---|---|---|---|
| SyNOnly_CC_2_reg_iter_1_bleached_masked | 4.833 | 1 | voldiff | 4.5 | 1 |
| SyNOnly_CC_2_reg_iter_1_bleached_masked | 5.5 | 3 | landmarks | 4.5 | 1 |
| SyNOnly_CC_2_reg_iter_1_bleached_masked | 4.333 | 2 | dice | 4.5 | 1 |
| SyNOnly_CC_2_reg_iter_1_normal_masked | 6.333 | 4 | voldiff | 6 | 2 |

| | | | | | |
|---|---|---|---|---|---|
| SyNOnly_CC_2_reg_iter_1_normal_masked | 4.75 | 2 | landmarks | 6 | 2 |
| SyNOnly_CC_2_reg_iter_1_normal_masked | 5.5 | 4 | dice | 6 | 2 |
| SyNOnly_CC_4_reg_iter_1_bleached_masked | 5.5 | 3 | voldiff | 6 | 2 |
| SyNOnly_CC_4_reg_iter_1_bleached_masked | 6 | 4 | landmarks | 6 | 2 |
| SyNOnly_CC_4_reg_iter_1_bleached_masked | 4.333 | 1 | dice | 6 | 2 |
| SyNOnly_CC_4_reg_iter_1_normal_masked | 10.167 | 10 | voldiff | 9.5 | 4 |
| SyNOnly_CC_4_reg_iter_1_normal_masked | 4.688 | 1 | landmarks | 9.5 | 4 |
| SyNOnly_CC_4_reg_iter_1_normal_masked | 8.5 | 7 | dice | 9.5 | 4 |
| SyNOnly_CC_6_reg_iter_1_bleached_masked | 7.667 | 5 | voldiff | 10 | 5 |
| SyNOnly_CC_6_reg_iter_1_bleached_masked | 6.938 | 6 | landmarks | 10 | 5 |
| SyNOnly_CC_6_reg_iter_1_bleached_masked | 5.333 | 3 | dice | 10 | 5 |
| SyNOnly_CC_2_reg_iter_0_bleached_masked | 5.333 | 2 | voldiff | 11 | 6 |
| SyNOnly_CC_2_reg_iter_0_bleached_masked | 8.25 | 7 | landmarks | 11 | 6 |
| SyNOnly_CC_2_reg_iter_0_bleached_masked | 7.5 | 6 | dice | 11 | 6 |

| | | | | | |
|---|---|---|---|---|---|
| SyNOnly_CC_6_reg_iter_1_normal_masked | 11.333 | 12 | voldiff | 15 | 7 |
| SyNOnly_CC_6_reg_iter_1_normal_masked | 6 | 4 | landmarks | 15 | 7 |
| SyNOnly_CC_6_reg_iter_1_normal_masked | 9.667 | 10 | dice | 15 | 7 |
| SyNOnly_MI_32_reg_iter_1_bleached_masked | 9 | 6 | voldiff | 16 | 8 |
| SyNOnly_MI_32_reg_iter_1_bleached_masked | 10.5 | 11 | landmarks | 16 | 8 |
| SyNOnly_MI_32_reg_iter_1_bleached_masked | 5.5 | 4 | dice | 16 | 8 |
| SyNOnly_CC_4_reg_iter_0_bleached_masked | 9.167 | 7 | voldiff | 16.5 | 9 |
| SyNOnly_CC_4_reg_iter_0_bleached_masked | 9.875 | 9 | landmarks | 16.5 | 9 |
| SyNOnly_CC_4_reg_iter_0_bleached_masked | 8.833 | 8 | dice | 16.5 | 9 |
| SyNOnly_CC_2_reg_iter_0_normal_masked | 9.833 | 8 | voldiff | 17.5 | 10 |
| SyNOnly_CC_2_reg_iter_0_normal_masked | 9.312 | 8 | landmarks | 17.5 | 10 |
| SyNOnly_CC_2_reg_iter_0_normal_masked | 10 | 11 | dice | 17.5 | 10 |
| SyNOnly_CC_6_reg_iter_0_bleached_masked | 9.833 | 8 | voldiff | 22.5 | 11 |
| SyNOnly_CC_6_reg_iter_0_bleached_masked | 11.062 | 13 | landmarks | 22.5 | 11 |

| | | | | | |
|---|---|---|---|---|---|
| SyNOnly_CC_6_reg_iter_0_bleached_masked | 10 | 11 | dice | 22.5 | 11 |
| SyNOnly_MI_32_reg_iter_1_normal_masked | 11.833 | 13 | voldiff | 23 | 12 |
| SyNOnly_MI_32_reg_iter_1_normal_masked | 10.438 | 10 | landmarks | 23 | 12 |
| SyNOnly_MI_32_reg_iter_1_normal_masked | 10.333 | 13 | dice | 23 | 12 |
| SyNOnly_CC_4_reg_iter_0_normal_masked | 11 | 11 | voldiff | 24.5 | 13 |
| SyNOnly_CC_4_reg_iter_0_normal_masked | 10.625 | 12 | landmarks | 24.5 | 13 |
| SyNOnly_CC_4_reg_iter_0_normal_masked | 12.833 | 14 | dice | 24.5 | 13 |
| SyNOnly_MI_32_reg_iter_0_bleached_masked | 11.833 | 13 | voldiff | 26 | 14 |
| SyNOnly_MI_32_reg_iter_0_bleached_masked | 12.625 | 15 | landmarks | 26 | 14 |
| SyNOnly_MI_32_reg_iter_0_bleached_masked | 9.167 | 9 | dice | 26 | 14 |
| SyNOnly_CC_6_reg_iter_0_normal_masked | 12.167 | 15 | voldiff | 28.5 | 15 |
| SyNOnly_CC_6_reg_iter_0_normal_masked | 11.062 | 13 | landmarks | 28.5 | 15 |
| SyNOnly_CC_6_reg_iter_0_normal_masked | 14.667 | 16 | dice | 28.5 | 15 |
| SyNOnly_MI_32_reg_iter_0_normal_masked | 13.833 | 17 | voldiff | 32 | 16 |

| | | | | | |
|---|---|---|---|---|---|
| SyNOnly_MI_32_reg_iter_0_normal_masked | 14.062 | 16 | landmarks | 32 | 16 |
| SyNOnly_MI_32_reg_iter_0_normal_masked | 13.167 | 15 | dice | 32 | 16 |
| TVMSQC_bleached_masked | 13 | 16 | voldiff | 34.5 | 17 |
| TVMSQC_bleached_masked | 17.188 | 18 | landmarks | 34.5 | 17 |
| TVMSQC_bleached_masked | 16.667 | 17 | dice | 34.5 | 17 |
| TVMSQ_bleached_masked | 15.333 | 18 | voldiff | 35 | 18 |
| TVMSQ_bleached_masked | 16.438 | 17 | landmarks | 35 | 18 |
| TVMSQ_bleached_masked | 16.667 | 18 | dice | 35 | 18 |
| TVMSQ_normal_masked | 15.833 | 19 | voldiff | 38 | 19 |
| TVMSQ_normal_masked | 17.312 | 19 | landmarks | 38 | 19 |
| TVMSQ_normal_masked | 17.833 | 19 | dice | 38 | 19 |
| TVMSQC_normal_masked | 16.167 | 20 | voldiff | 40 | 20 |
| TVMSQC_normal_masked | 17.375 | 20 | landmarks | 40 | 20 |
| TVMSQC_normal_masked | 19.167 | 20 | dice | 40 | 20 |

**Supplementary Table 9: Ranking nonlinear registration settings neonatal to template**

| Registration setting tested | meanRanks | consensusrank | metric | summed_consesnsus_ranking | final_ranking |
|---|---|---|---|---|---|
| SyNOnly_CC_2_reg_iter_1_normal_masked | 1.167 | 1 | dice | 3 | 1 |
| SyNOnly_CC_2_reg_iter_1_normal_masked | 3.667 | 1 | voldiff | 3 | 1 |
| SyNOnly_CC_2_reg_iter_1_normal_masked | 2.375 | 2 | landmarks | 3 | 1 |
| SyNOnly_CC_4_reg_iter_1_normal_masked | 2.333 | 2 | dice | 3 | 1 |
| SyNOnly_CC_4_reg_iter_1_normal_masked | 4.333 | 2 | voldiff | 3 | 1 |
| SyNOnly_CC_4_reg_iter_1_normal_masked | 2.188 | 1 | landmarks | 3 | 1 |
| SyNOnly_CC_6_reg_iter_1_normal_masked | 3.5 | 3 | dice | 5.5 | 3 |
| SyNOnly_CC_6_reg_iter_1_normal_masked | 4.333 | 2 | voldiff | 5.5 | 3 |
| SyNOnly_CC_6_reg_iter_1_normal_masked | 3 | 3 | landmarks | 5.5 | 3 |
| SyNOnly_MI_32_reg_iter_1_normal_masked | 3.667 | 4 | dice | 8 | 4 |
| SyNOnly_MI_32_reg_iter_1_normal_masked | 4.833 | 4 | voldiff | 8 | 4 |
| SyNOnly_MI_32_reg_iter_1_normal_masked | 4.625 | 4 | landmarks | 8 | 4 |
| SyNOnly_CC_2_reg_iter_0_normal_masked | 4.5 | 5 | dice | 10 | 5 |

| | | | | | |
|---|---|---|---|---|---|
| SyNOnly_CC_2_reg_iter_0_normal_masked | 5.333 | 5 | voldiff | 10 | 5 |
| SyNOnly_CC_2_reg_iter_0_normal_masked | 4.812 | 5 | landmarks | 10 | 5 |
| SyNOnly_CC_4_reg_iter_0_normal_masked | 6.333 | 6 | dice | 12.5 | 6 |
| SyNOnly_CC_4_reg_iter_0_normal_masked | 5.5 | 7 | voldiff | 12.5 | 6 |
| SyNOnly_CC_4_reg_iter_0_normal_masked | 6.062 | 6 | landmarks | 12.5 | 6 |
| SyNOnly_CC_6_reg_iter_0_normal_masked | 7.333 | 8 | dice | 13.5 | 7 |
| SyNOnly_CC_6_reg_iter_0_normal_masked | 5.333 | 5 | voldiff | 13.5 | 7 |
| SyNOnly_CC_6_reg_iter_0_normal_masked | 6.25 | 7 | landmarks | 13.5 | 7 |
| SyNOnly_MI_32_reg_iter_0_normal_masked | 7.167 | 7 | dice | 15.5 | 8 |
| SyNOnly_MI_32_reg_iter_0_normal_masked | 6.333 | 8 | voldiff | 15.5 | 8 |
| SyNOnly_MI_32_reg_iter_0_normal_masked | 7.312 | 8 | landmarks | 15.5 | 8 |
| TVMSQ_normal_masked | 9.167 | 9 | dice | 17.5 | 9 |
| TVMSQ_normal_masked | 6.333 | 8 | voldiff | 17.5 | 9 |
| TVMSQ_normal_masked | 8.938 | 9 | landmarks | 17.5 | 9 |

| | | | | | |
|---|---|---|---|---|---|
| TVMSQC_normal_masked | 9.833 | 10 | dice | 20 | 10 |
| TVMSQC_normal_masked | 9 | 10 | voldiff | 20 | 10 |
| TVMSQC_normal_masked | 9.438 | 10 | landmarks | 20 | 10 |

**Supplementary Table 10: Ranking nonlinear registration settings fetal to neonatal**

| algorithm | meanRanks | consensusrank | metric | summed_consesnsus_ranking | final_ranking |
|---|---|---|---|---|---|
| SyNOnly_CC_2_reg_iter_1_bleached_masked | 3.5 | 1 | landmarks | 2.5 | 1 |
| SyNOnly_CC_2_reg_iter_1_bleached_masked | 2.333 | 1 | dice | 2.5 | 1 |
| SyNOnly_CC_2_reg_iter_1_bleached_masked | 6.667 | 2 | voldiff | 2.5 | 1 |
| SyNOnly_CC_4_reg_iter_1_bleached_masked | 5.75 | 3 | landmarks | 5 | 2 |
| SyNOnly_CC_4_reg_iter_1_bleached_masked | 2.333 | 1 | dice | 5 | 2 |
| SyNOnly_CC_4_reg_iter_1_bleached_masked | 7.167 | 3 | voldiff | 5 | 2 |
| SyNOnly_CC_2_reg_iter_0_bleached_masked | 6.438 | 4 | landmarks | 6.5 | 3 |
| SyNOnly_CC_2_reg_iter_0_bleached_masked | 4.667 | 4 | dice | 6.5 | 3 |
| SyNOnly_CC_2_reg_iter_0_bleached_masked | 6.5 | 1 | voldiff | 6.5 | 3 |

| | | | | | |
|---|---|---|---|---|---|
| SyNOnly_CC_2_reg_iter_1_normal_masked | 5.25 | 2 | landmarks | 10 | 4 |
| SyNOnly_CC_2_reg_iter_1_normal_masked | 6.833 | 6 | dice | 10 | 4 |
| SyNOnly_CC_2_reg_iter_1_normal_masked | 8.833 | 10 | voldiff | 10 | 4 |
| SyNOnly_CC_6_reg_iter_1_bleached_masked | 7.625 | 5 | landmarks | 10.5 | 5 |
| SyNOnly_CC_6_reg_iter_1_bleached_masked | 3.667 | 3 | dice | 10.5 | 5 |
| SyNOnly_CC_6_reg_iter_1_bleached_masked | 8.333 | 8 | voldiff | 10.5 | 5 |
| SyNOnly_CC_4_reg_iter_0_bleached_masked | 7.875 | 6 | landmarks | 11 | 6 |
| SyNOnly_CC_4_reg_iter_0_bleached_masked | 6.833 | 6 | dice | 11 | 6 |
| SyNOnly_CC_4_reg_iter_0_bleached_masked | 7.667 | 4 | voldiff | 11 | 6 |
| SyNOnly_MI_32_reg_iter_1_bleached_masked | 8.625 | 8 | landmarks | 12.5 | 7 |
| SyNOnly_MI_32_reg_iter_1_bleached_masked | 6.167 | 5 | dice | 12.5 | 7 |
| SyNOnly_MI_32_reg_iter_1_bleached_masked | 7.667 | 4 | voldiff | 12.5 | 7 |
| SyNOnly_CC_4_reg_iter_1_normal_masked | 8.125 | 7 | landmarks | 18 | 10 |
| SyNOnly_CC_4_reg_iter_1_normal_masked | 9.5 | 10 | dice | 18 | 10 |

| | | | | | |
|---|---|---|---|---|---|
| SyNOnly_CC_4_reg_iter_1_normal_masked | 11.5 | 12 | voldiff | 18 | 10 |
| SyNOnly_CC_2_reg_iter_0_normal_masked | 8.875 | 9 | landmarks | 17.5 | 8 |
| SyNOnly_CC_2_reg_iter_0_normal_masked | 9.167 | 9 | dice | 17.5 | 8 |
| SyNOnly_CC_2_reg_iter_0_normal_masked | 8.333 | 8 | voldiff | 17.5 | 8 |
| SyNOnly_CC_6_reg_iter_0_bleached_masked | 9.312 | 10 | landmarks | 17.5 | 8 |
| SyNOnly_CC_6_reg_iter_0_bleached_masked | 7.667 | 8 | dice | 17.5 | 8 |
| SyNOnly_CC_6_reg_iter_0_bleached_masked | 8.167 | 7 | voldiff | 17.5 | 8 |
| SyNOnly_MI_32_reg_iter_0_bleached_masked | 9.438 | 11 | landmarks | 19 | 11 |
| SyNOnly_MI_32_reg_iter_0_bleached_masked | 9.5 | 10 | dice | 19 | 11 |
| SyNOnly_MI_32_reg_iter_0_bleached_masked | 8 | 6 | voldiff | 19 | 11 |
| SyNOnly_CC_6_reg_iter_1_normal_masked | 10.25 | 12 | landmarks | 24.5 | 12 |
| SyNOnly_CC_6_reg_iter_1_normal_masked | 11.667 | 12 | dice | 24.5 | 12 |
| SyNOnly_CC_6_reg_iter_1_normal_masked | 12.5 | 13 | voldiff | 24.5 | 12 |
| SyNOnly_CC_4_reg_iter_0_normal_masked | 11.062 | 13 | landmarks | 25 | 13 |

| | | | | | |
|---|---|---|---|---|---|
| SyNOnly_CC_4_reg_iter_0_normal_masked | 12.667 | 13 | dice | 25 | 13 |
| SyNOnly_CC_4_reg_iter_0_normal_masked | 11 | 11 | voldiff | 25 | 13 |
| SyNOnly_MI_32_reg_iter_1_normal_masked | 12.125 | 14 | landmarks | 27.5 | 14 |
| SyNOnly_MI_32_reg_iter_1_normal_masked | 12.833 | 14 | dice | 27.5 | 14 |
| SyNOnly_MI_32_reg_iter_1_normal_masked | 12.5 | 13 | voldiff | 27.5 | 14 |
| SyNOnly_CC_6_reg_iter_0_normal_masked | 12.375 | 15 | landmarks | 30 | 15 |
| SyNOnly_CC_6_reg_iter_0_normal_masked | 14.5 | 15 | dice | 30 | 15 |
| SyNOnly_CC_6_reg_iter_0_normal_masked | 12.667 | 15 | voldiff | 30 | 15 |
| SyNOnly_MI_32_reg_iter_0_normal_masked | 14.688 | 16 | landmarks | 32 | 16 |
| SyNOnly_MI_32_reg_iter_0_normal_masked | 15.667 | 16 | dice | 32 | 16 |
| SyNOnly_MI_32_reg_iter_0_normal_masked | 13.833 | 16 | voldiff | 32 | 16 |
| TVMSQ_bleached_masked | 16.5 | 17 | landmarks | 34.5 | 17 |
| TVMSQ_bleached_masked | 17.5 | 17 | dice | 34.5 | 17 |
| TVMSQ_bleached_masked | 14.333 | 18 | voldiff | 34.5 | 17 |

| | | | | | |
|---|---|---|---|---|---|
| TVMSQC_normal_masked | 17.375 | 19 | landmarks | 36 | 18 |
| TVMSQC_normal_masked | 18.5 | 18 | dice | 36 | 18 |
| TVMSQC_normal_masked | 13.833 | 16 | voldiff | 36 | 18 |
| TVMSQC_bleached_masked | 17.25 | 18 | landmarks | 37.5 | 19 |
| TVMSQC_bleached_masked | 19.5 | 20 | dice | 37.5 | 19 |
| TVMSQC_bleached_masked | 15.167 | 19 | voldiff | 37.5 | 19 |
| TVMSQ_normal_masked | 17.562 | 20 | landmarks | 39 | 20 |
| TVMSQ_normal_masked | 18.5 | 18 | dice | 39 | 20 |
| TVMSQ_normal_masked | 15.333 | 20 | voldiff | 39 | 20 |

#### References

Avants, B. B., C. L. Epstein, M. Grossman, and J. C. Gee. 2008. “Symmetric Diffeomorphic Image Registration with Cross-Correlation: Evaluating Automated Labeling of Elderly and Neurodegenerative Brain.” *Medical Image Analysis* 12 (1): 26–41.

Cox, R. W. 1996. “AFNI: Software for Analysis and Visualization of Functional Magnetic Resonance Neuroimages.” *Computers and Biomedical Research, an International Journal* 29 (3): 162–73.

Jenkinson, Mark, Peter Bannister, Michael Brady, and Stephen Smith. 2002. “Improved Optimization for the Robust and Accurate Linear Registration and Motion Correction of Brain Images.” *NeuroImage* 17 (2): 825–41.

Jenkinson, Mark, Christian F. Beckmann, Timothy E. J. Behrens, Mark W. Woolrich, and Stephen M. Smith. 2012. “FSL.” *NeuroImage* 62 (2): 782–90.

Tustison, Nicholas J., Philip A. Cook, Andrew J. Holbrook, Hans J. Johnson, John Muschelli, Gabriel A. Devenyi, Jeffrey T. Duda, et al. 2021. “The ANTsX Ecosystem for Quantitative Biological and Medical Imaging.” *Scientific Reports* 11 (1): 9068.
